# Genomic, Epigenomic, and Transcriptional Characterisation of Carbapenem and Colistin Resistance Mechanisms in *Klebsiella pneumoniae* and *Enterobacter* species

**DOI:** 10.1101/2023.12.15.571804

**Authors:** Masego Mmatli, Nontombi Marylucy Mbelle, P. Bernard Fourie, John Osei Sekyere

## Abstract

The emergence of colistin and carbapenem-resistant *Klebsiella pneumoniae* isolates presents a significant global health threat. This study investigates the resistance mechanisms in six *K. pneumoniae* and four *Enterobacter* sp. isolates lacking carbapenemases or *mcr* genes using genomics and transcriptomics. The ten isolates were classified into three categories: non-carbapenemase-producing, carbapenem-resistant strains (n = 4), non-*mcr*–producing colistin-resistant strains (n = 5), and one isolate susceptible to both antibiotics.

The analysis included phenotypic characterization using MicroScan ID/AST, enzyme (MCR and Metallo β-lactamase) and efflux pump inhibition (EPI) assays. Whole-genome sequencing, RNA sequencing, and bioinformatics tools were employed in subsequent analysis. Most of the *K. pneumoniae* were ST307 with KL102 and O1/O2V2 serotypes. MicroScan revealed multidrug resistance, and AMR analysis identified numerous ARGs in *K. pneumoniae*. *Enterobacter* species possessed fewer resistance genes; nevertheless, they encoded virulence factors and gene mutations, potentially impacting the AST profile. *K. pneumoniae* ARGs were mainly plasmid-borne, with IncFIB(K)/IncFII(K) in Kp_15 harbouring up to nineteen ARGs. Virulence factors included biofilm formation, capsule production, and type IV secretion. Epigenomic investigations revealed prevalent type I (*M1.Ecl34977I*) and type II (*M.Kpn34618Dcm*) restriction modification sites. Compared to international isolates, the study isolates phylogenetically clustered more closely with Chinese strains. Transcriptomics showed high efflux pump activity in carbapenem-resistant isolates, confirmed by EPI. Further, mutations were identified in outer membrane proteins. Colistin-resistant isolates exhibited high capsule production, efflux pump, and putative glycotransferase activity, potentially influencing their phenotypes.

In conclusion, genomic and transcriptional analyses enhanced our understanding of adaptive mechanisms in clinical multidrug-resistant pathogens, posing significant public health challenges.

## Introduction

*Klebsiella pneumoniae*, a member of the Enterobacteriaceae family, is frequently identified as the aetiological agent of infections caused by carbapenem-resistant bacteria worldwide ^1^. Infections caused by *K. pneumoniae* include urinary and respiratory tract infections as well as bloodstream infections in neonates ^2^.

Management of *K. pneumoniae* infections has resulted in the overuse of antibiotics and the emergence and rapid dissemination of super bugs resistant to both carbapenems and colistin ^3^. Carbapenem-resistant *K. pneumoniae* (CRKP) in the clinical setting is largely mediated by the acquisition of carbapenemases, which are commonly associated with mobile genetic elements (MGEs). These MGEs include plasmids, transposons and integrons, ^4^ which facilitate wide resistance gene dissemination between animal- and human pathogens ^5^. In South Africa, there have been several reports of carbapenemase-producing *K. pneumoniae* outbreaks in the clinical setting ^6-9^. Carbapenemases that have been identified in South Africa include *Klebsiella pneumoniae* carbapenemase (KPC), Verona Integron-Mediated Metallo-β-lactamase (VIM), Imipenemase (IMP), New Delhi metallo β-lactamase (NDM), and oxacillinase (OXA) ^4^. Amongst these carbapenemases, *bla*_OXA_ and *bla*_NDM_ genes are the most common and primarily reported in South Africa ^4^.

*Bla_OXA_*_-181_-producing *K. pneumoniae* have caused several outbreaks in several provinces in South Africa, with the ST307 being the most predominant clone ^4,6,8-10^. Other carbapenem-resistance mechanisms include decreased membrane permeability through increased efflux activity and decreased porin expression; these are usually coupled with β-lactamase activity^11^. An observational study performed in the United States found that carbapenemase-producing Enterobacteriaceae (CPE) infections have an increased risk of fatality than non-CPE infections^12^, thus highlighting the health risk imposed by these microorganisms ^12^.

Colistin is the last-resort antibiotic that is currently being used, interchangeably, with tigecycline to manage CRKP isolates. Unfortunately, there is a high prevalence of colistin resistance in CRKP clinical isolates ^13^. Although not common in South African clinical settings, *mcr* genes are responsible for majority of colistin resistance in Enterobacteriaceae, particularly in *Escherichia coli* ^7,14,15^. The inactivation of *mgrB*, which inhibits the kinase activity of *PhoPQ,* is the most common colistin resistance mechanism in *K. pneumoniae* ^16,17^. The two-component system (TCS), PhoPQ, are regulators of the *pbgP* operon that encodes the endogenous lipopolysaccharide modification system. This operon is also regulated by the PmrAB TCS. Thus, mutations within *phoP, phoQ, pmrA* and *pmrB* results in the modification of the LPS, ^18^ which reduces the negative net charge of the LPS ^11,19,20^.

Other colistin resistance mechanisms include the use of efflux pumps, the formation of capsules and decreasing the outer membrane proteins ^11^. The prevalence of colistin- and carbapenem-resistant *K. pneumoniae* is increasing in South Africa and globally, necessitating surveillance studies that will monitor their epidemiology and resistance mechanisms. ^21^

This study aims to characterize novel colistin and carbapenem resistance mechanisms in six clinical *K. pneumoniae* isolates and four *Enterobacter* sp isolates using both genomics and RNA-seq. These clinical isolates were part of a molecular screening that evaluated the epidemiology of carbapenemases and *mcr* genes in Pretoria, South Africa ^7^.

## Methods

### 2.1 Study Settings and Samples Collection

The ten clinical isolates identified by MicroScan to be *K. pneumoniae* were obtained from a collection of multi-drug resistant (MDR) Gram-negative bacteria during a molecular screening study ^7^. These isolates were collected from the National Health Laboratory Service, Tshwane Academic Division (NHLS/TAD), a referral laboratory. At the time of collection, the clinical isolates were classified as carbapenem and/or colistin resistant at collection. They were specifically selected because they tested negative for known carbapenemases and *mcr ge*nes, including *bla*_IMP_, *bla*_KPC_, *bla*_NDM_, *bla*_OXA-48_, *bla*_NDM_, *bla*_VIM_ and *mcr 1-5* genes, as determined by multi-plex PCR screening^7^. Ethics approval for this study was obtained from the Faculty of Health Sciences Research Ethics Committee of the University of Pretoria (Ref no. 581/2020).

### 2.2 Phenotypic testing

#### 2.2.1 Minimum inhibitory concentration evaluation

The ten clinical isolates presumed to be *K. pneumoniae* were cultured on blood agar plates and incubated at 37□ for 24 hours. After incubation, the isolates underwent antimicrobial susceptibility testing and species identification using MicroScan automated system with Combo 66 panels (Beckman Coulter). The results were interpreted according to the Clinical and Laboratory Standard Institute (CLSI) guidelines ^22^.

For the carbapenem- and colistin-resistant isolates, a manual broth microdilution assay was performed following ISO standard 20776-1 ^23^. Ertapenem sulphate salt and colistin sulphate salt (Glentham Life Sciences, United Kingdom), were used for the assay ^24^. *E. coli* ATCC 25922 was included as a quality control strain. Both antibiotics were dissolved in sterile deionized water according to the manufacturers’ instructions. The antibiotic concentrations tested were: 128 µg/mL, 64 µg/mL, 32 µg/mL, 16 µg/mL, 8 µg/mL, 4 µg/mL, 2 µg/mL, 1 µg/mL, 0.5 µg/mL, and 0.25 µg/mL.

The assay was performed in untreated 96-well polystyrene microtiter plates, with each well containing 100 µL of antibiotic dilution and Mueller-Hinton broth (MHB) or cation-adjusted MHB for ertapenem and colistin respectively. Subsequently, a 0.5 MacFarland suspension of bacterial strains was prepared, diluted it 1:20 with sterile saline, and added 0.01 mL of bacterial inoculum to each well. The plates also included sensitive and negative control wells.

Following inoculation, the plates were incubated at 37 °C for 16-18 hours, and the minimum inhibitory concentration (MIC) was determined as the lowest antibiotic concentration without visible bacterial growth ^22^. Its important to note that since the completion of this study, CLSI revised their colistin resistance breakpoint to ≥ 4 mg/mL, rendering the previous breakpoint of ≥ 2 mg/mL used in this study outdated and incorrect.

#### 2.2.2 Conditional treatment with carbapenems and colistin

Conditional treatment was performed on the ten *K. pneumoniae* isolates before RNA extraction. The carbapenem-resistant isolates were exposed to 0.5 mg/mL of ertapenem, while the colistin-resistant isolates were exposed to 2 mg/mL of colistin. Briefly, 1 mL of a 0.5 M *K. pneumoniae* suspension was transferred to 2 mL Eppendorf tubes, and the appropriate volumes of antibiotics were added to achieve final concentrations of 0.5 mg/mL for ertapenem and 2 mg/mL for colistin. The sensitive isolate served as a control and was left untreated. Subsequently, all ten isolates were incubated at 37°C for 16-18 hours.

#### 2.2.3 Treatment with efflux pump inhibitors and EDTA

To evaluate the change in susceptibility of ertapenem and colistin in the presence of an efflux pump inhibitor (EPIs) and EDTA, the same procedure described above in the “MIC Evaluation” section was followed. The EPIs used were carbonyl cyanide m-chlorophenylhydrazone (CCCP), reserpine (RES), verapamil (VER), and phenylalanine-arginine β-naphthylamide (PaβN). The EPIs CCCP, PaβN, and RES were diluted in dimethyl sulfoxide (DMSO), while VER was diluted in sterile distilled water.

The final concentrations of the substrates in the broth were 1.5 µg/mL for CCCP, 4 µg/mL for VER, 25 µg/mL for PAβN, 20 µg/mL for RES, and 20 mM (pH 8.0) for EDTA. Efflux pump, Metallo β-lactamase, and MCR activity were determined by observing a 2-fold or greater reduction in MICs of ertapenem and colistin.

### 2.3 Molecular Investigations of Resistance Mechanisms

#### 2.3.1 Nucleic acid extraction

For nucleic acid extractions, fresh pure colonies grown on Mueller-Hinton Agar (Diagnostic Media Products) were used. DNA and RNA were extracted using commercial kits: Quick-DNA-fungal/bacterial MiniPrep™ kit (ZymoResearch) was used for DNA and Quick-RNA-fungal/bacterial MiniPrep™ kit (Zymo Research) was used for RNA. The extraction protocols followed the manufacturers’ instructions, and the concentration and purity of the DNA extracts were checked using the NanoDrop™ 2000/2000c Spectrophotometer (Thermo Fisher Scientific Inc.) before sequencing. RNA samples were stored at -80°C, while the DNA samples were stored at -20°C until sequencing.

#### 2.3.2 Whole-genome sequencing and RNA-sequencing

The extracted DNA samples were sent to the National Institute of Communicable Diseases (NICD) Sequencing Core Facility for whole genome sequencing using PacBio SMRT sequencing at 100x coverage. The RNA samples were sent to Inqaba Biotechnology for PacBio Isoform sequencing, which provides long and accurate HiFi reads for a diverse transcriptome.

#### 2.3.3 Genomic analysis

The sequenced genomes were submitted to Genbank and assigned accession numbers under the Bioproject PRJNA861833. The Centre for Genomic Epidemiology pipeline (http://www.genomicepidemiology.org/services/) was used to analyse the sequenced DNA and retrieve information about the species identity, multi locus sequence type (MLST), antibiotic resistance genes (ARGs), and plasmids harboured by each sequenced isolate. The Kaptive-web database (https://kaptive-web.erc.monash.edu/) was used to predict the *K. pneumoniae* isolates’ serotypes (K types and O types). VRprofile2 platform (https://tool2-mml.sjtu.edu.cn/VRprofile/home.php) was used to associate ARGs and virulence genes to their mobilome. PacBio’s hierarchical genome-assembly process (HGAP) software was used to assemble the PacBio reads Spades was used to assemble the Illumina reads.

#### 2.3.4 Epigenomic analyses

The restriction modification system (RMS), which includes DNA methylation, restriction endonucleases, and their motifs, was identified for each isolate using the Restriction Enzyme Database (REBASE), hosted by the Centre for Epidemiology. The PacBio MotifMaker software was used for determining methylation modifications and motifs. Owing to financial constraints, this analysis was only conducted on three *K. pneumoniae* isolates (Kp_14, Kp_24, and H3) and two *Enterobacter* sp. isolates (A5 and G5), which were selected for PacBio SMRT sequencing.

#### 2.3.5 Phylogenetics

The genetic relationships among *Enterobacter* sp. isolates, specifically focusing on *E. cloacae*, *E. bugandensis*, and *E. asburiae* was investigated. For each species, three phylogenetic trees were generated using global whole genome sequences of *Enterobacter* sp. Each tree included genomes of the respective species, including *E. cloacae* (n = 33), *E. bugandensi*s (n = 26), and *E. asburiae* (n = 53).

In the case of *K. pneumoniae* isolates, a phylogenetic reconstruction was performed using 82 whole genome sequences obtained from various settings, including South Africa (n = 28), other African regions (n = 11), and globally (n = 43). This analysis aimed to assess the epidemiological and evolutionary links between the clinical *K. pneumoniae* isolates examined in this study and other *K. pneumoniae* species within these three distinct geographical settings.

The 194 whole genome sequences used in the phylogenetic analysis were retrieved from the PATRIC website (https://www.bv-brc.org/), and comprehensive data on these strains are provided in Table S1. *Escherichia coli* ATCC 25922 (Genbank accession number: CP009073) served as the reference genome. The phylogenetic analysis was conducted using PATRIC’s phylogenetic tree building service, which employs the randomized axelerated maximum likelihood (RAxML) program.

### 2.4 RNA-sequencing data analysis

The RNA-sequencing data analysis was conducted using the HTSeq-DeSeq2 tool for aligning, assembling, and evaluating the differential expression data from the different sample groups. Each *K. pneumoniae* isolate was compared with the carbapenem- and colistin-susceptible strain, Kp13; *K. pneumoniae* MGH64 was used as the reference genome. The differentially expressed genes (DEGs) were identified using the *K. pneumoniae* strain MGH64 genome. The function of each gene was evaluated using the genome annotations of the reference strain on the PATRIC platform.

## Results

### 3.1 Strain description

Ten putative *K. pneumoniae* isolates were selected from a collection of 302 clinical MDR Gram-negative bacteria during a molecular screening study of carbapenemases and *mcr* genes^7^. These ten isolates included a carbapenem- and colistin-sensitive strain and were categorized into three groups. The first group comprised of four strains that did not produce carbapenemases but were resistant to carbapenems. The second group consisted of isolates resistant to colistin without producing *mcr* genes. Specifically, the carbapenem-resistant isolates were Kp_4, Kp_14, Kp_15, and Kp_24, while the colistin-resistant ones were A3, G3, G5, G8 and H3. As detailed in the method section, these isolates were exposed to ertapenem and colistin for RNA-seq. The third group was the sensitive strain, Kp_13, which displayed susceptibility to both colistin and ertapenem, and served as a reference genome for the subsequent RNA-seq.

### 3.2 Phenotypic characterization

#### 3.2.1 MIC and MicroScan analysis

The ten isolates underwent Microscan analysis using the Neg Combo 66 panel for identification and antimicrobial susceptibility testing of 25 antibiotics, including ertapenem, imipenem, meropenem, and colistin. Table 1 reveals that seven isolates had an MIC > 2 µg/mL indicating resistance to colistin, while three isolates, Kp_4, Kp_13, and Kp_15 showed susceptibility to colistin with an MIC value of ≤ 2. Among the non-*mcr-*producing isolates (A5, G3, G5, G8, and H3), colistin MIC values greater than 4 µg/mL were observed. The BMD assay (using ertapenem) demonstrated that these isolates had an MIC value of 128 µg/mL while *E. coli* ATCC 25922 had an MIC value of 0.25 µg/mL (Table 2).

**Table 1.**
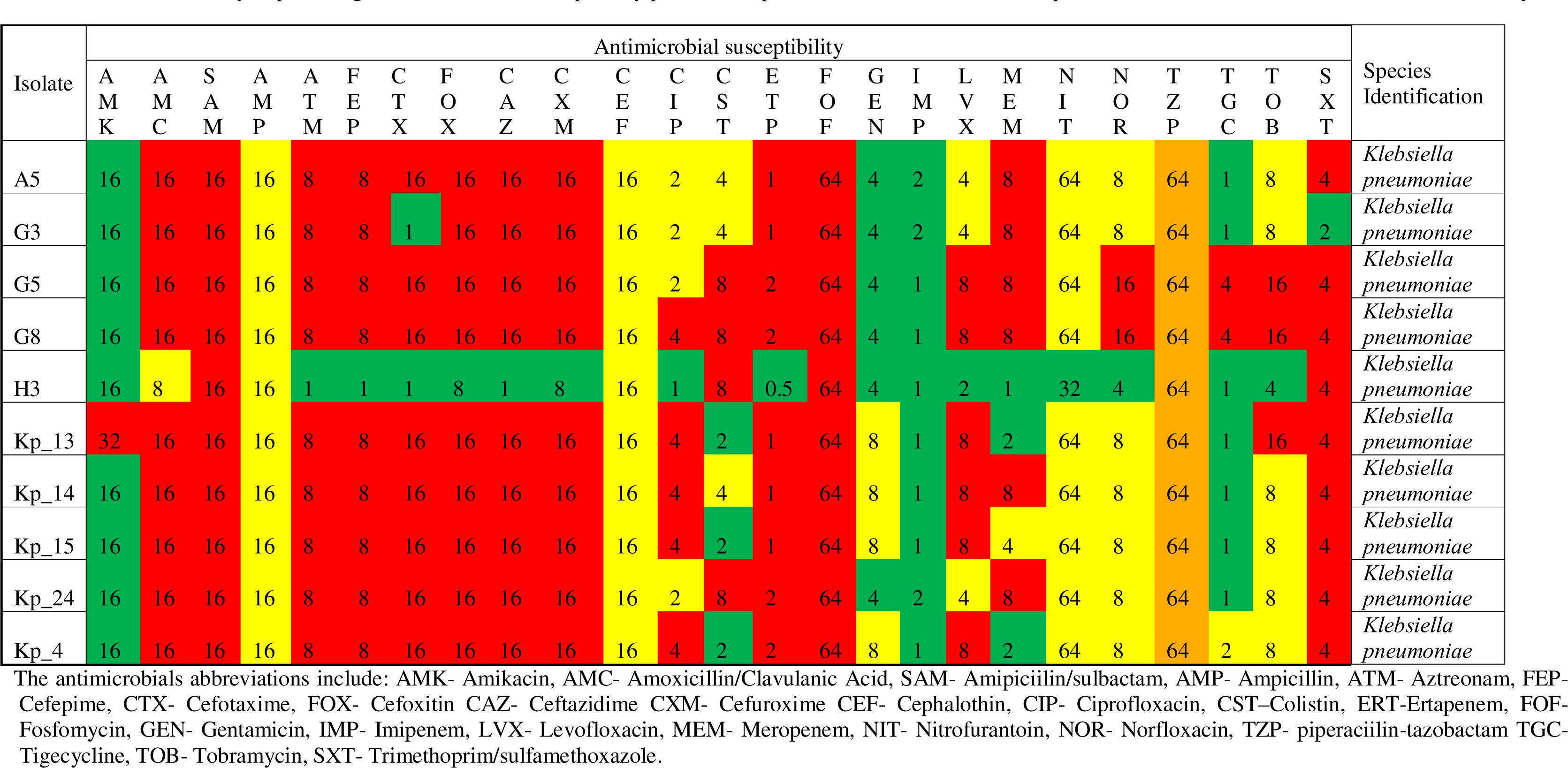
MicroScan analysis providing the antimicrobial susceptibility profile and species identification of the carbapenem and colistin resistant isolates included in study.

| Isolate | Antimicrobial susceptibility |  |  |  |  |  |  |  |  |  |  |  |  |  |  |  |  |  |  |  |  |  |  |  |  | Species Identification |
| --- | --- | --- | --- | --- | --- | --- | --- | --- | --- | --- | --- | --- | --- | --- | --- | --- | --- | --- | --- | --- | --- | --- | --- | --- | --- | --- |
|  | A<br>M<br>K | A<br>M<br>C | S<br>A<br>M | A<br>M<br>P | A<br>T<br>M | F<br>E<br>P | C<br>T<br>X | F<br>O<br>X | C<br>A<br>Z | C<br>X<br>M | C<br>E<br>F | C<br>I<br>P | C<br>S<br>T | E<br>R<br>T | F<br>O<br>F | G<br>E<br>N | I<br>M<br>P | L<br>V<br>X | M<br>E<br>M | N<br>I<br>T | N<br>O<br>R | T<br>Z<br>P | T<br>G<br>C | T<br>O<br>B | S<br>X<br>T |  |
| A5 | 16 | 16 | 16 | 16 | 8 | 8 | 16 | 16 | 16 | 16 | 16 | 2 | 4 | 1 | 64 | 4 | 2 | 4 | 8 | 64 | 8 | 64 | 1 | 8 | 4 | Klebsiella pneumoniae |
| G3 | 16 | 16 | 16 | 16 | 8 | 8 | 1 | 16 | 16 | 16 | 16 | 2 | 4 | 1 | 64 | 4 | 2 | 4 | 8 | 64 | 8 | 64 | 1 | 8 | 2 | Klebsiella pneumoniae |
| G5 | 16 | 16 | 16 | 16 | 8 | 8 | 16 | 16 | 16 | 16 | 16 | 2 | 8 | 2 | 64 | 4 | 1 | 8 | 8 | 64 | 16 | 64 | 4 | 16 | 4 | Klebsiella pneumoniae |
| G8 | 16 | 16 | 16 | 16 | 8 | 8 | 16 | 16 | 16 | 16 | 16 | 4 | 8 | 2 | 64 | 4 | 1 | 8 | 8 | 64 | 16 | 64 | 4 | 16 | 4 | Klebsiella pneumoniae |
| H3 | 16 | 8 | 16 | 16 | 1 | 1 | 1 | 8 | 1 | 8 | 16 | 1 | 8 | 0.5 | 64 | 4 | 1 | 2 | 1 | 32 | 4 | 64 | 1 | 4 | 4 | Klebsiella pneumoniae |
| Kp_13 | 32 | 16 | 16 | 16 | 8 | 8 | 16 | 16 | 16 | 16 | 16 | 4 | 2 | 1 | 64 | 8 | 1 | 8 | 2 | 64 | 8 | 64 | 1 | 16 | 4 | Klebsiella pneumoniae |
| Kp_14 | 16 | 16 | 16 | 16 | 8 | 8 | 16 | 16 | 16 | 16 | 16 | 4 | 4 | 1 | 64 | 8 | 1 | 8 | 8 | 64 | 8 | 64 | 1 | 8 | 4 | Klebsiella pneumoniae |
| Kp_15 | 16 | 16 | 16 | 16 | 8 | 8 | 16 | 16 | 16 | 16 | 16 | 4 | 2 | 1 | 64 | 8 | 1 | 8 | 4 | 64 | 8 | 64 | 1 | 8 | 4 | Klebsiella pneumoniae |
| Kp_24 | 16 | 16 | 16 | 16 | 8 | 8 | 16 | 16 | 16 | 16 | 16 | 2 | 8 | 2 | 64 | 4 | 2 | 4 | 8 | 64 | 8 | 64 | 1 | 8 | 4 | Klebsiella pneumoniae |
| Kp_4 | 16 | 16 | 16 | 16 | 8 | 8 | 16 | 16 | 16 | 16 | 16 | 4 | 2 | 2 | 64 | 8 | 1 | 8 | 2 | 64 | 8 | 64 | 2 | 8 | 4 | Klebsiella pneumoniae |

The antimicrobials abbreviations include: AMK- Amikacin, AMC- Amoxicillin/Clavulanic Acid, SAM- Amipiciilin/sulbactam, AMP- Ampicillin, ATM- Aztreonam, FEP-Cefepime, CTX- Cefotaxime, FOX- Cefoxitin CAZ- Ceftazidime CXM- Cefuroxime CEF- Cephalothin, CIP- Ciprofloxacin, CST–Colistin, ERT-Ertapenem, FOF-Fosfomycin, GEN- Gentamicin, IMP- Imipenem, LVX- Levofloxacin, MEM- Meropenem, NIT- Nitrofurantoin, NOR- Norfloxacin, TZP- piperaciilin-tazobactam TGC-Tigecycline, TOB- Tobramycin, SXT- Trimethoprim/sulfamethoxazole.

**Table 2.** Broth Microdilution assay evaluating the effect of EDTA and EPIs on the MIC value (µg/mL) of Ertapenem and Colistin ERT: ertapenem, CST: colistin, PaβN: phenylalanine-arginine β-naphthylamide, CCCP: carbonyl cyanide m-chlorophenylhydrazone, RES: Reserpine, VER: Verapamil, EPIs: Efflux pump inhibitors

| Isolate | ERT-MIC | ERT-MIC in presence of EPIs/EDTA |  |  |  |  |
| --- | --- | --- | --- | --- | --- | --- |
|  |  | EDTA | PAβN | CCCP | RES | VER |
| Kp_4 | 16 | 4 | 16 | 8 | 16 | 16 |
| Kp_13 | 0.5 | 0.5 | 0.5 | 0.5 | 0.5 | 0.5 |
| Kp_14 | 16 | 2 | 16 | 16 | 8 | 16 |
| Kp_15 | 64 | 32 | 64 | 32 | 32 | 32 |
| Kp_24 | 128 | 128 | 128 | 128 | 128 | 128 |
| *E. coli* ATCC 25922 | 0.25 | 0.25 | 0.25 | 0.25 | 0.25 | 0.25 |
| *P. aeruginosa* ATCC 27853 | 4 | 4 | 4 | 4 | 4 | 4 |
| Isolate | CST-MIC | CST-MIC in presence of EPIs/EDTA |  |  |  |  |
|  |  | EDTA | PAβN | CCCP | RES | VER |
| A5 | 128 | 128 | 128 | 128 | 128 | 128 |
| G3 | 128 | 128 | 128 | 128 | 128 | 128 |
| G5 | 128 | 128 | 128 | 64 | 128 | 128 |
| G8 | 128 | 128 | 128 | 64 | 128 | 128 |
| H3 | 128 | 128 | 128 | 64 | 128 | 128 |
| *E. coli* ATCC 25922 | 0.25 | 0.25 | 0.25 | 0.25 | 0.25 | 0.25 |
| *P. aeruginosa* ATCC 27853 | 0.25 | 0.25 | 0.25 | 0.25 | 0.25 | 0.25 |

From the Microscan analysis, nine isolates were resistant to ertapenem (MIC > 0.5 µg/mL) while all the ten isolates were susceptible to imipenem (MIC ≤ 2 µg/mL) (Table 1). Additionally, seven isolates were resistant to meropenem (MIC > 2 µg/mL) (Table 1). The non-carbapenemase-producing isolates viz., Kp_4, Kp_14, Kp_15, and Kp_24, were resistant to ertapenem (MIC > 2 µg/mL) but were susceptible to imipenem (MICs ≤ 2 µg/mL).

Finally, all isolates, except Kp_4 (MIC of 2 µg/mL), displayed non-susceptibility to meropenem (MIC > 2 µg/mL). The isolates included in the study were MDR isolates, three of which were non-susceptible to tigecycline (Table 1). Kp_13 was susceptible to colistin, imipenem, and meropenem: MICs of 2, 1, and 2 µg/mL, respectively.

The MicroScan analysis identified all isolates as *K. pneumoniae* (Table 1).

#### 3.2.2 Effects of EDTA and EPIs on MIC values of ertapenem and colistin

The addition of EDTA significantly impacted the ertapenem MICs of Kp_4, Kp_14, and Kp_15 isolates, while no growth inhibition was observed in Kp_24 (Table 2). Furthermore, CCCP reduced the ertapenem MIC values of Kp_4 and Kp_15 with the MIC of Kp_4 decreasing from 16 µg/ml to 8 µg/ml and the MIC of Kp_15 decreasing from 64 µg/ml to 32 µg/ml. Additionally, RES decreased the ertapenem MIC value of Kp_15 from 16 µg/ml to 8 µg/ml. However, no growth inhibition was observed in Kp_24 with the addition of EPIs.

In non-*mcr-*producing colistin-resistant isolates, the effects of EDTA and EPIs were evaluated (Table 2). The addition of EDTA did not inhibit the growth of the isolates in the presence of colistin. However, a decrease in MIC values was observed when CCCP was added to G5, G8 and H3, with their colistin MIC values decreasing from 128 µg/ml to 64 µg/ml. No growth inhibition was observed for the other EPIs tested.

### 3.3 Genomic characterization

The whole-genome sequencing analysis identified six isolates as *K. pneumoniae*, the remaining isolates were two *Enterobacter cloacae* complex strains, one *Enterobacter asburiae* and one *Enterobacter bugandensis* isolate (Table 3). Among the *K. pneumoniae* isolates, four MLST groups were identified: ST307 (Kp_4, Kp_15 and Kp_24), ST219 (Kp_14), ST25 (H3), and a novel sequence type, ST6408, for Kp_13.

**Table 3.** Genomic identification and characterization of the 10 presumed Klebsiella pneumoniae isolates included in the study.

| Isolate | Species | Serotypes | MLST | Plasmids | Antibiotic resistance genes |  |
| --- | --- | --- | --- | --- | --- | --- |
|  |  |  |  |  | Chromosomal | Plasmids |
| Kp4 | *Klebsiella pneumoniae* | K: KL102<br>O: O1/O2v2 | ST307 | IncFIA(HI1) IncFIB(K)/IncFII(K) IncL IncR | blaSHV-28, fosA6, oqxA, oqxB | blaCTX-M-15, blaTEM-1B, aac(3)-Iia, aac(6')-Ib-cr, aadA16, aph(3'')-Ib, aph(6)-Id, ARR-3, dfrA27, qacE, qnrB6, sul1, sul2, tetD |
| Kp13 | *Klebsiella pneumoniae* | K: KL142<br>O: O1/O2v1 | ST6408 | IncFIA(HI1)/IncR/repB(R1701) IncFIB(K)/IncFIB(K)/IncR | blaSHV-81, fosA6, oqxA, oqxB | blaCTX-M-14, blaDHA-1, blaTEM-1B, aac(3)-Iid, aac(6')-Ib-cr, aadA16, aph(3')-Ia, aph(3'')-Ib, aph(6)-Id, armA, ARR-3, dfrA27, floR, mphA, mphE, msrE, qacE, qnrB4, sul1, sul2, tetA |
| Kp14 | *Klebsiella pneumoniae* | K: KL114<br>O: O1/O2v1 | ST219 | IncFIB(K)(pCAV1099-114) | blaSHV-26, fosA | blaCTX-M-15, aadA2, aph(3')-Ia, aph(3'')-Ib, aph(6)-Id, catA2, dfrA12, mphA, qacE, qnrS1, sul1, sul2 |
| Kp15 | *Klebsiella pneumoniae* | K: KL102<br>O: O1/O2v2 | ST307 | IncFIB(K)(pCAV1099-114) IncFIB(K)/IncFII(K) IncX3 | blaSHV-28, fosA6, oqxA, oqxB | blaCTX-M-15, blaOXA-1, blaOXA-181, blaTEM-1B, aac(3)-Iia, aac(6)-Ib-cr, aadA2, aph(3')-Ia, aph(3'')-Ib, aph(6)-Id, catA2, catB3, dfrA12, dfrA14, mphA, qacE, qnrB1, qnrS1, sul1, sul2, tetA |
| Kp24 | *Klebsiella pneumoniae* | K: KL102<br>O: O1/O2v2 | ST307 | IncFIB(pNDM-Mar)/IncHI1B(pNDM-MAR) IncX3 | blaSHV-28, fosA6, oqxA, oqxB | blaCTX-M-15, blaOXA-1, blaOXA-181, blaTEM-1C, aac(6)-Ib-cr, aadA1, ant(3'')-Ia, catB3, dfrA15, mphA, qacE, qnrS1, sul1 |
| H3 | *Klebsiella pneumoniae* | K: KL2<br>O: O1/O2v1 | ST25 | IncFIB(K)/IncFII(K) | blaCMH-3, blaSHV-81, fosA, fosA6, oqxA, oqxB | blaTEM-1B, aph(3')-Ia, aph(3'')-Ib, aph(6)-Id, dfrA14, mphA, sul2 |
| A5 | *Enterobacter asburiae* | N/A | ST22 | None | blaACT-4, fosA, oqxA, oqxB | None |
| G3 | *Enterobacter bugandensis* | N/A | ST632 | None | blaACT-6, fosA, oqxA, oqxB | None |
| G5 | *Enterobacter cloacae* | N/A | ST2100 | None | blaCMH-3, fosA, oqxA, oqxB | None |
| G8 | *Enterobacter cloacae* | N/A | ST2100 | None | blaCMH-3, fosA, oqxA, oqxB | None |

The analysis of K-loci and O-loci serotype revealed that the ST307 isolates (Kp4, Kp15 and Kp24) shared the same KL102 and O1/O2v2 results. The remaining isolates all had the same O1/O2v2 O-loci type. However, KL142, KL114 and KL2 K-loci types were found in Kp13, Kp14 and H3, respectively (Table 3).

Twelve plasmids were identified within the six *K. pneumoniae* isolates. These plasmids were associated with ten compatibility groups, with IncFIB(K), IncFII(K), and IncR being the most common. Eight of these plasmids co-harboured multiple compatibility groups, while the remaining four were singletons (Tables 3 and S2). Among the isolates, Kp_4 hosted the highest number of plasmids (n = 4), followed by Kp_15 (n=3). Isolates Kp_13 and Kp_25 each carried two plasmids, while both Kp_4 and H3 only hosted one plasmid.

The largest plasmid observed belonged to Kp_15, with a size of 311.9 kbp. This plasmid consisted of two incompatibility groups, namely IncFII(K) and IncFIB(K). The second largest plasmid belonged to H3, with a size of 216.8 kbp. This plasmid consisted of multiple replicons, including IncFIB(K), IncFII(K), and IncQ1. Notably, no plasmids were identified within the *Enterobacter* sp. isolates.

### 3.4 Antibiotic resistance gene analysis

All the isolates harboured β-lactamase genes that influenced their phenotypic β-lactam resistance, corroborating the PCR results from the molecular screening (Table 3). ^7^ The *Enterobacter* species (A5 and G3) harboured β-lactamase genes, namely *bla_ACT-6_* within the chromosome, while G5 and G8 harboured *bla*_CMH-3_ genes (Table 3 and Table S2). These β-lactamase genes were not found in association with mobile genetic elements (MGEs). The *K. pneumoniae* isolates harboured multiple β-lactamase genes. Notably, *bla*_SHV_ variants, which are intrinsic to *K. pneumoniae,* were found in H3, Kp_13, Kp_14 and Kp_4, along with *bla*_CMH-3_ genes, all of which were located within the chromosome. Isolate H3 additionally harboured *bla*_TEM-1B,_ another β-lactamase gene, located on an unidentified plasmid (Table S2). Kp_13 isolate harboured four additional β-lactamase genes including *bla*_CTX-M-15_ and *bla*_TEM-1B,_ which were surrounded by MGEs *IS*26 and *IS*Kpn26, respectively (Table S2).

Two other genes, *bla*_DHA-1_ and *bla*_TEM-1B,_ were located on the IncFIB(K) plasmid and surrounded by *IS*26 and *IS*Kpn26, respectively. Kp_14 harboured four additional *bla*_CTX-M-15_ genes located on three contigs, along with chromosomal *bla*_SHV-26._ Two of the *bla*_CTX-M-15_ genes were harboured on an IncFIB plasmid, while the other two were situated on an unidentified plasmid or transposable elements. Kp_15 harboured four additional β-lactamase, including chromosomal *bla*_SHV-28_ and IncFIB(K)/IncFII(K) plasmid-borne *bla*_CTX-M-15_, *bla*_OXA-1,_ and *bla*_TEM-1B._ Additionally, *bla*_OXA-181,_ was located on the IncX3 plasmid, also surrounded by IS26. Kp_24 harboured four additional β-lactamase genes, including *bla*_SHV-28;_ *bla*_OXA-181_ was located on an IncX3 plasmid, also surrounded by *IS*26. The remaining genes, *bla*_OXA-1_, *bla*_CTX-M-15,_ and *bla*_TEM-1B_, were located on an unidentified plasmid or transposable element, and were all surrounded by *IS*26.

Lastly, Kp_4 harboured five additional β-lactamase genes, including chromosomal *bla*_SHV-28,_ two *bla*_CTX-M-15,_ and three *bla*_TEM-1B_. Two *bla*_TEM-1B_ and one *bla*_CTX-M-15_ were located on separate unidentified plasmids or transposable elements, while the remaining *bla*_TEM-1B_ and *bla*_CTX-M-15_ were located on the IncFIA(HI1) plasmid (Table S2).

The four *Enterobacter* species (A5, G3, G5, and G8) harboured β-lactamase genes and three additional chromosomal antibiotic resistance genes (ARGs): *fosA, oqxA,* and *oqxB*. These ARGs were also present within the chromosomes of the *K. pneumoniae* isolates, Kp_4, Kp_13, Kp_14, Kp_24, and H3 (Table 3 and Table S3). However, the resistance genes *oqxA* and *oqxB* were not found in isolate Kp_14. The remaining ARGs listed in Table 3 were located on plasmids or extrachromosomal DNA, and included genes mediating resistance to aminoglycosides (*aac(3’)-IIa, acc(6’)-Ib-cr, aadA1, aadA16, aadA2, ant(3”)-Ia, aph(3’)-Ia, aph(3”)-Ib, aph(6)-Id, armA)*, amphenicol (*catA2/B3, floR*), macrolide (*mphE, mphA, msrE*), quaternary ammonium compound (*qacE*), quinolone (*qnrB1/B4/B6/S1*), sulphonamide (*sul1, sul2*), tetracycline (*tetA, tetD*), and trimethoprim (*dfrA12/14/15/27*). The pathogen watch pipeline identified *ompK35* mutations conferring carbapenem resistance in isolates Kp_4, Kp_13, Kp_15, Kp_24, and H3 (Table S3). However, the pipeline failed to analyse the *Enterobacter* species isolates. Isolate H3 was further found to harbour *mgrB* mutations conferring resistance to colistin (Table S3).

### 3.5 Virulence genes analysis

Thirty virulence genes were identified on chromosomes within the ten isolates (Table S4); they were flanked by MGEs. On average, each isolate carried ten virulence genes, with G8 harbouring the lowest of four genes, and Kp_14 harbouring the highest number of 19 virulence genes. Certain virulence genes were found within prophage MGEs including *algU* (present in H3 and Kp_15), *hcp/tssD* (Kp_14, Kp_15, Kp_24, and Kp_4), and *rfaE* (H3). Additionally, the *hcp/tssD* gene found in Kp_4 was located near integrative conjugative elements within the chromosome.

The different categories of virulence genes include those responsible for biofilm formation, capsular synthesis, the type VI secretion system (T6SS), and lipopolysaccharide synthesis. Biofilm formation genes were only observed in isolate Kp_14. These genes include *fimA, fimC, fimD, fimF, fimG*, and *fimI*, which are responsible for type 1 fimbriae and are involved in biofilm formation. Several capsular synthesis virulence genes were identified within *K. pneumoniae*. These include *gnd* (Kp_4, Kp_13, Kp_14, Kp_15, and Kp_24), *manB/manC* (Kp_14), *ugd* (Kp_4, Kp_13, Kp_14, and Kp_24), *wcaJ* (Kp_13), and *wza* (Kp_13, Kp_14, Kp_15, and Kp_24).

Multiple lipopolysaccharide synthesis genes were identified. The following genes were identified in all six *K. pneumoniae* isolates: *glf, wbbM, wbbN*, and *wzt*. The remaining genes, *wzm* (Kp_4, Kp_13, Kp_14, and Kp_24), *wbtL* (Kp_13), *wbbO* (Kp_4, Kp_14, Kp_24, and H3), and *kfoC* (Kp_4, Kp_15, Kp_24, and H3), were only found in some *K. pneumoniae* isolates. Seven genes responsible for the T6SS were identified within both *K. pneumoniae* and *Enterobacter* species. The structural genes include *hcp/tssD* (A5, G3, H3, Kp_4, Kp_14, Kp_15, and Kp_24), *icmF/tssM* (A5, G3, G5, G8, Kp_13, and Kp_14), *sciN/tssJ* (A5 and G3), *tssF* (A5, G3, G5, G8), *tssG* (A5, G3, G5, G8), and lastly *tli1* (A5 and G3). Notably, A5 harboured all the structural genes. The last T6SS virulence gene identified as KPHS_23120, which was harboured by A5 and G3.

### 3.6 Phylogenetic analysis

#### 3.6.1 Phylogenetic analysis of the K. pneumoniae isolates

The phylogenetic analysis of the *K. pneumoniae* isolates included 81 isolates originating from five continents: Africa (n = 39), Asia (n = 15), Europe (n = 21), North America (n = 6), and South America (n = 4). These isolates belonged to nine sequence types (STs), with ST307 (n = 45), ST25 (n = 19), and ST219 (n = 12) being the most common clones. ST307 was found in eight countries, while ST25 and ST219 were found in seven countries. All *K. pneumoniae* isolates included in the phylogenetic analysis were obtained from human hosts.

The genome-based phylogeny of the South African *K. pneumoniae* isolates revealed six clades (Figure 1). Among the 28 *K. pneumoniae* isolates, 21 belonged to ST307, making up three of the six clades (Clades 4 to 6). These three clades had similar resistomes, with the highest similarities observed between Kp8, Tembi-19, Tembi-37, EC0361298, and EC03605938. In contrast, Clade 6 showed the least similarity within its isolates’ resistome.

**Figure 1.**
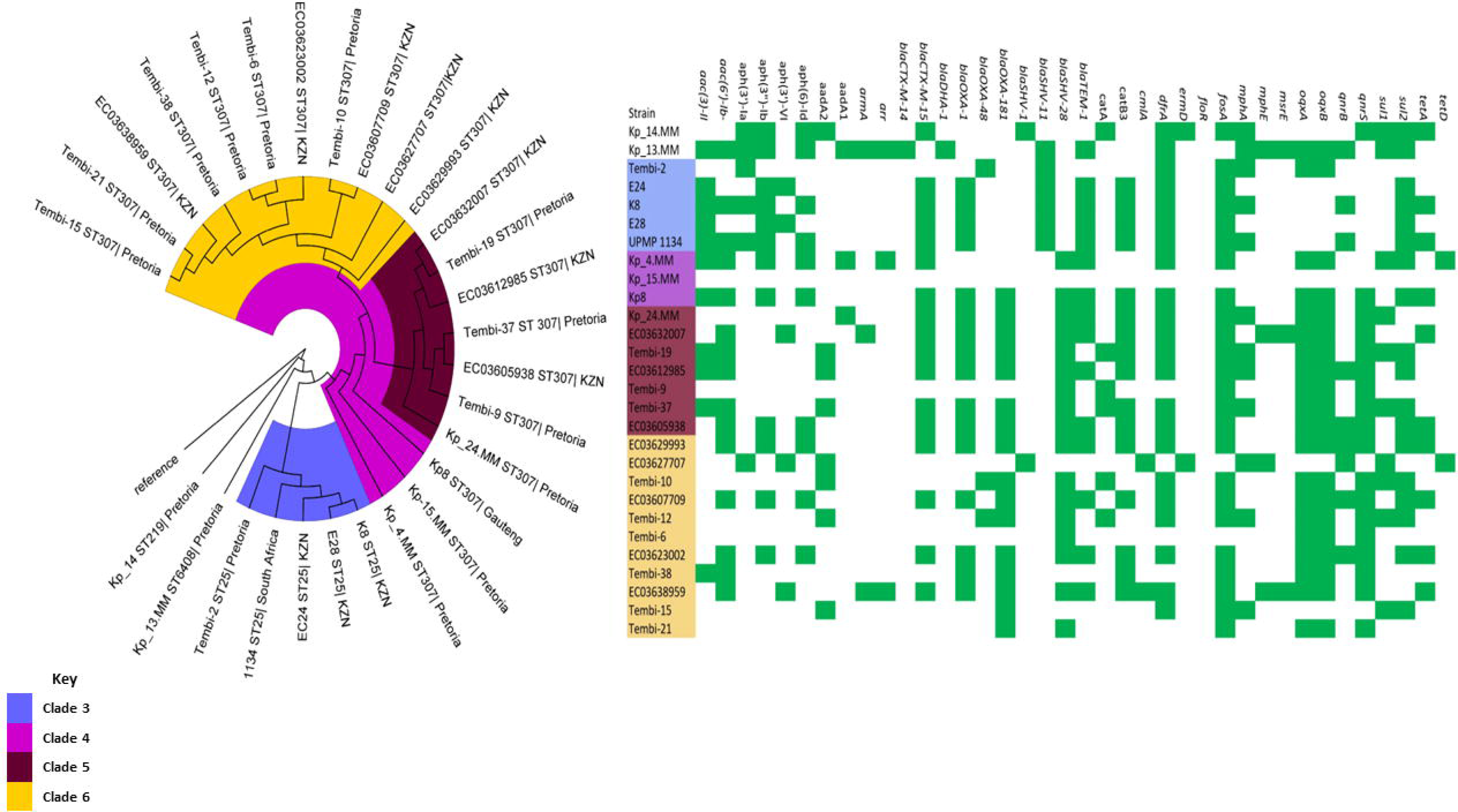
Phylogenetic and resistome dynamics of *K. pneumoniae* isolates from South Africa collected from human samples. Each strain is represented by its strain identifier, MLST designation, and country of origin. Strains belonging to the same clade are highlighted with the same color on the branches. The resistome is depicted through green and white blocks, representing the presence and absence of antibiotic resistance genes, respectively.

The phylogeny of the African *K. pneumoniae* isolates (Figure 2), consisted of seven clades with a high similarity within each clade concerning their resistomes. Clade 5 and 6 had similar resistome patterns. Interestingly, H2 ST501, which formed its own clade, shows its distinct resistome pattern, setting it apart from the other clades.

**Figure 2.**
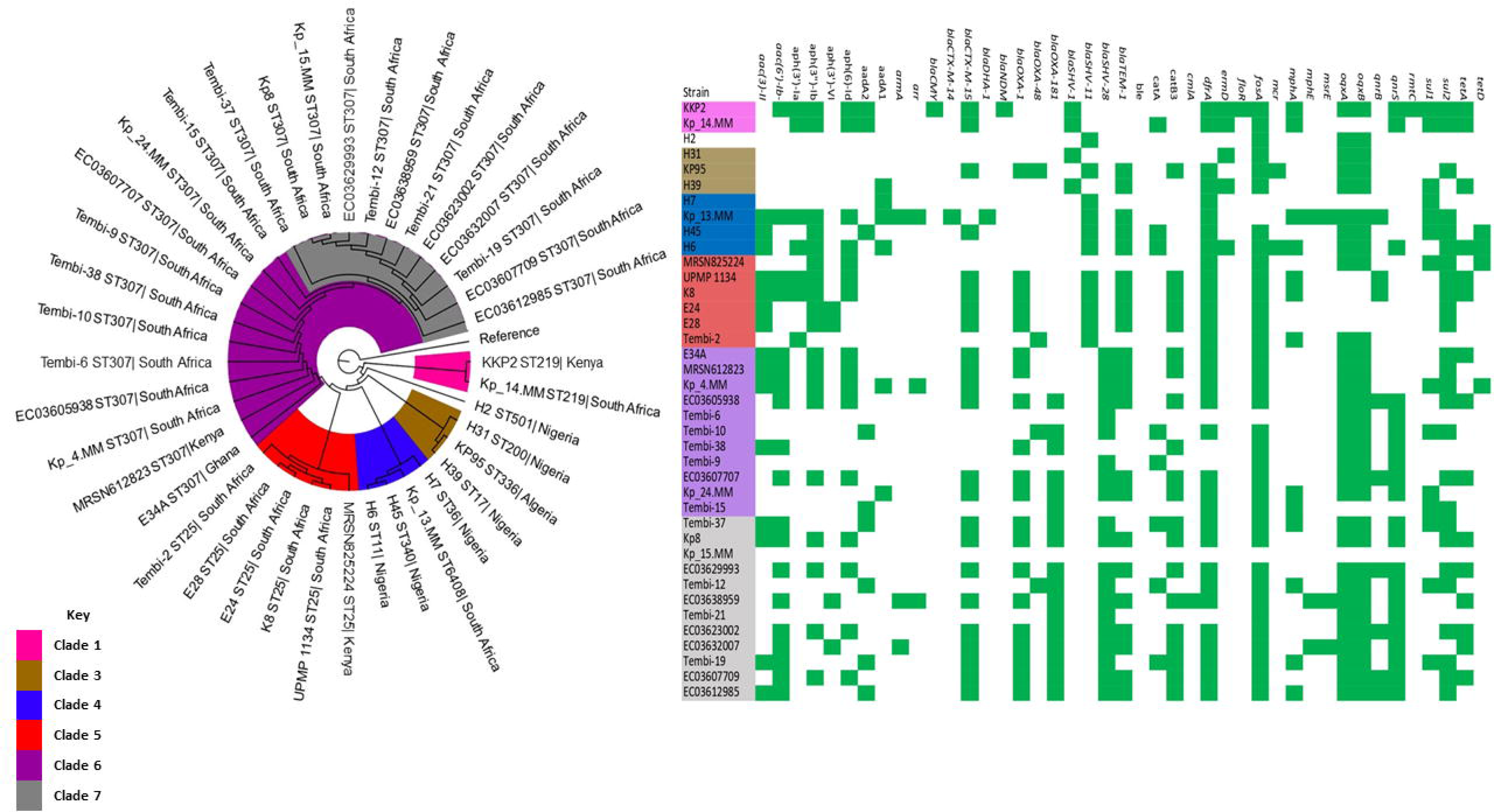
Phylogenetic and resistome dynamics of *K. pneumoniae* isolates from Africa collected from human samples. Each strain is represented by its strain identifier, MLST designation, and country of origin. Strains belonging to the same clade are highlighted with the same color on the branches. The resistome is depicted through green and white blocks, representing the presence and absence of antibiotic resistance genes, respectively.

Figure 3 shows the genome-based phylogeny of *K. pneumoniae* from the remaining continents, revealing six clades. Kp_14 was grouped in Clade 3 alongside other *K. pneumoniae* ST219 isolates and H2 ST501 from Nigeria. Kp_13 was placed in Clade 4, along with the three Nigerian *K. pneumoniae* isolates. Lastly, Kp_4, Kp_15, and Kp_24 were assigned to Clade 5 along with *K. pneumoniae* ST307 isolates.

**Figure 3.**
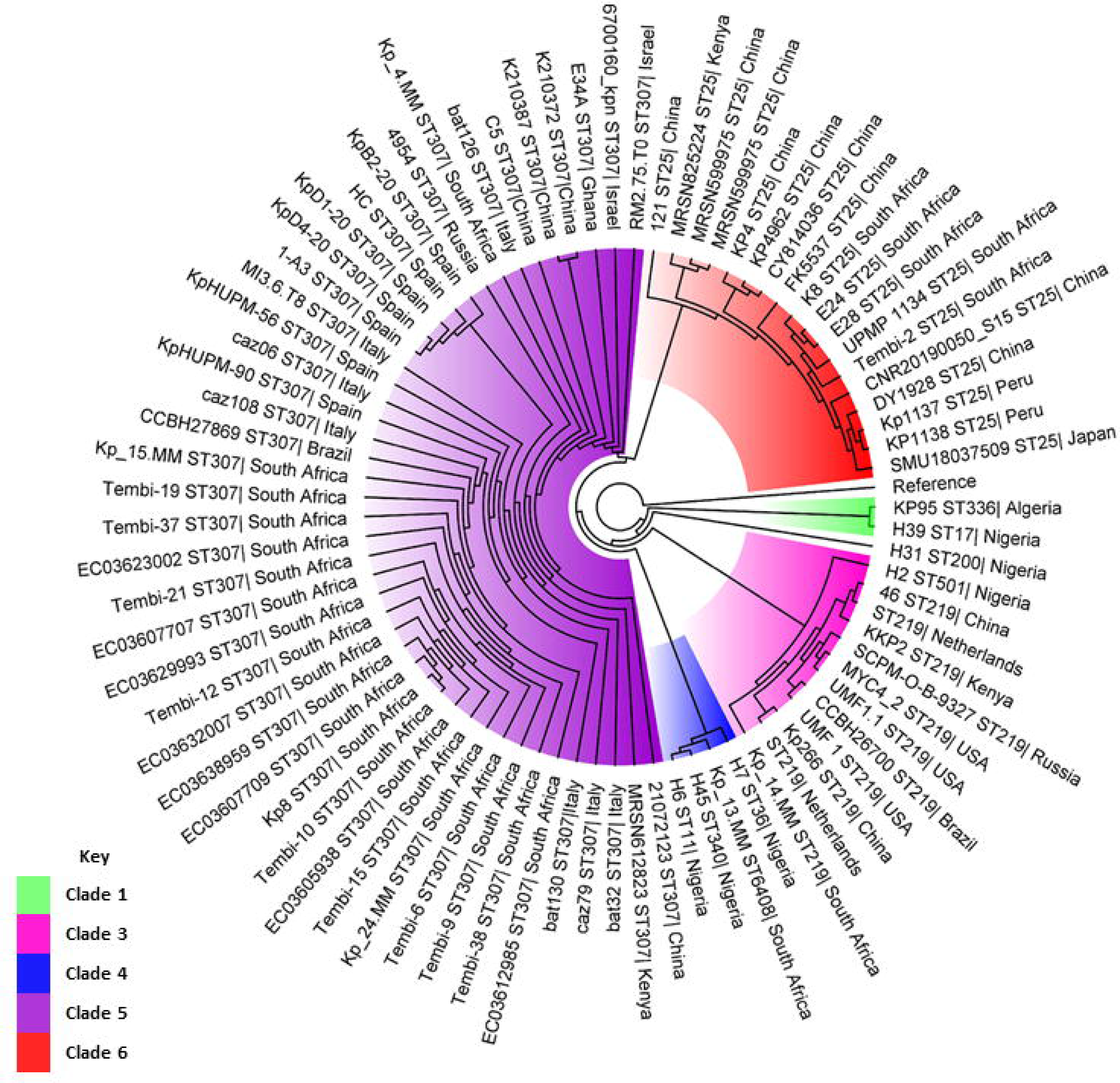
Global phylogenetic analysis of *K. pneumoniae* isolates collected from human samples. Each strain is represented by its strain identifier, MLST designation, and country of origin. Strains belonging to the same clade are highlighted with the same color on the branches.

#### 3.6.2 Phylogenetic analysis of the Enterobacter sp. isolates

For the *Enterobacter* species (*E. asburiae, E. bugandensis,* and *E. cloacae),* three separate phylogenetic trees were constructed. The phylogeny of *E. asburiae* seen in Figure 4, included 53 isolates distributed among seven distinct clades. Interestingly, isolate A5 was placed in clade 3 alongside a South African strain (E124_11) and a Chinese strain (C210176) forming a clade with a significantly similar resistome. Clades 6 and 7 harboured a wide range of ARGs, these two clades included isolates from six to seven countries, with China being the predominant source for both. In this phylogenetic tree, the clades exhibit the presence of *bla*_ACT_, fosA, and oqxB genes across most resistomes. Additionally, distinct resistome patterns are observed within each clade, indicating variations in the genes responsible for resistance mechanisms among the different groups.

**Figure 4.**
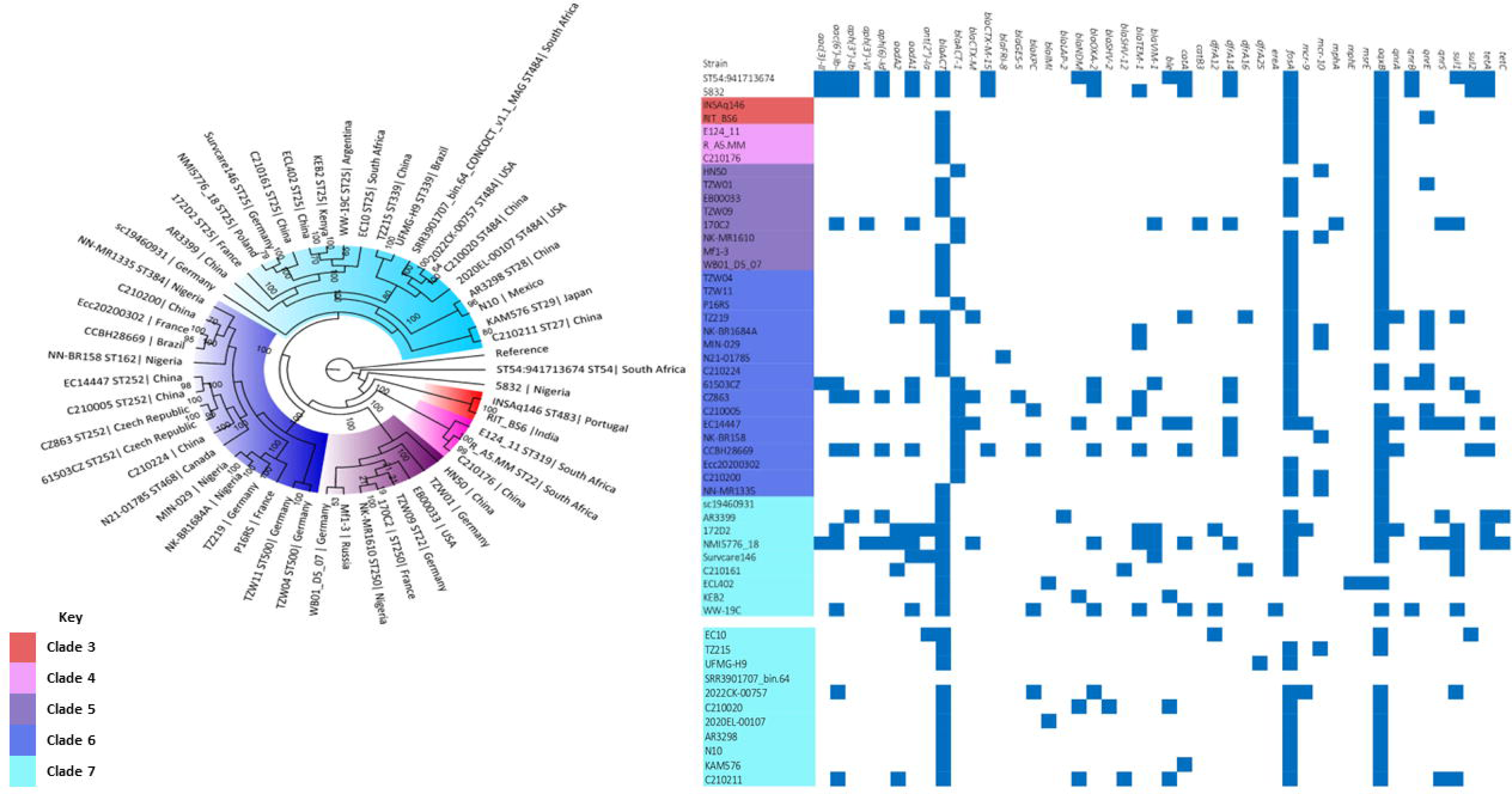
Global phylogenetic and resistome dynamics of *E. asburiae* isolates, collected from human samples. Each strain is represented by its strain identifier, MLST designation, and country of origin. Strains belonging to the same clade are highlighted with the same color on the branches. The resistome is depicted through blue and white blocks, representing the presence and absence of antibiotic resistance genes, respectively.

The genome phylogeny of *E. bugandensis* seen in Figure 5, included 25 isolates distributed among three distinct clades. The phylogenetic tree included three isolates that carried ten or more ARGs: IMP80 (Clade 1); C210207 and AR2787 (both in Clade 2). The remaining isolates harboured similar ARGs including *bla*_ACT_, found in all isolates, and *qnrA*, found in most isolates (n = 21). Compared with the other phylogenetic trees, this specific tree showed a lower number of resistance genes, with *bla*_ACT_ and *oqxB* being the predominant ARGs among the included isolates. Only four isolates harboured more than the average three ARGs. Excluding these isolates, a consistent and similar resistance pattern is observed across the tree, suggesting a commonality in resistance mechanisms acquired by *E. bugandensis* species.

**Figure 5.**
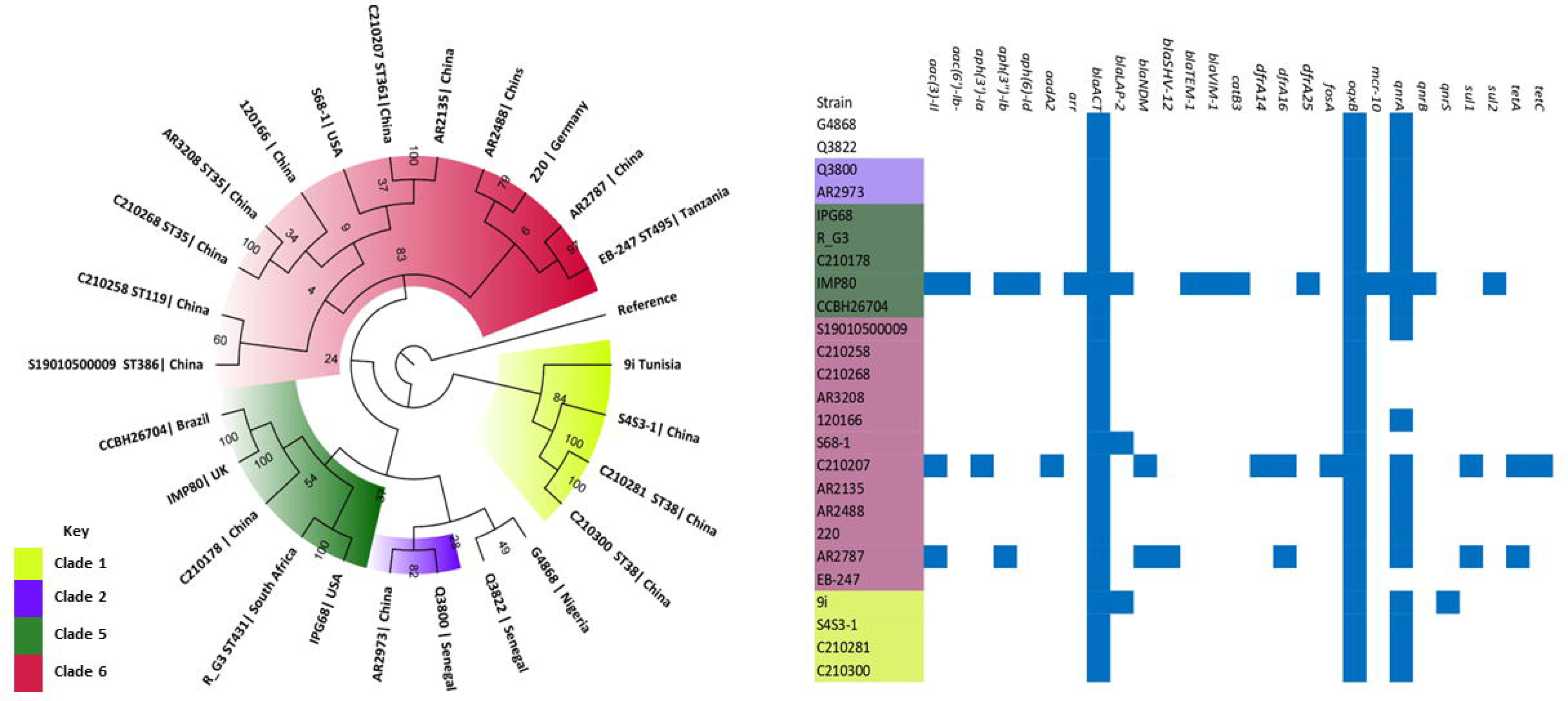
Global phylogenetic and resistome dynamics of *E. bugandensis* isolates, collected from human samples. Each strain is represented by its strain identifier, MLST designation, and country of origin. Strains belonging to the same clade are highlighted with the same color on the branches. The resistome is depicted through blue and white blocks, representing the presence and absence of antibiotic resistance genes, respectively.

The phylogeny of *E. cloacae* (Figure 6) included 32 isolates distributed among seven distinct clades. Clade 5 had the fewest ARGs followed by Clade 6, while Clade 2 and 4 harboured the most. All the isolates from Clade 4 originated from South Africa, while Clade 6 displayed a greater diversity in terms of countries of origin.

**Figure 6.**
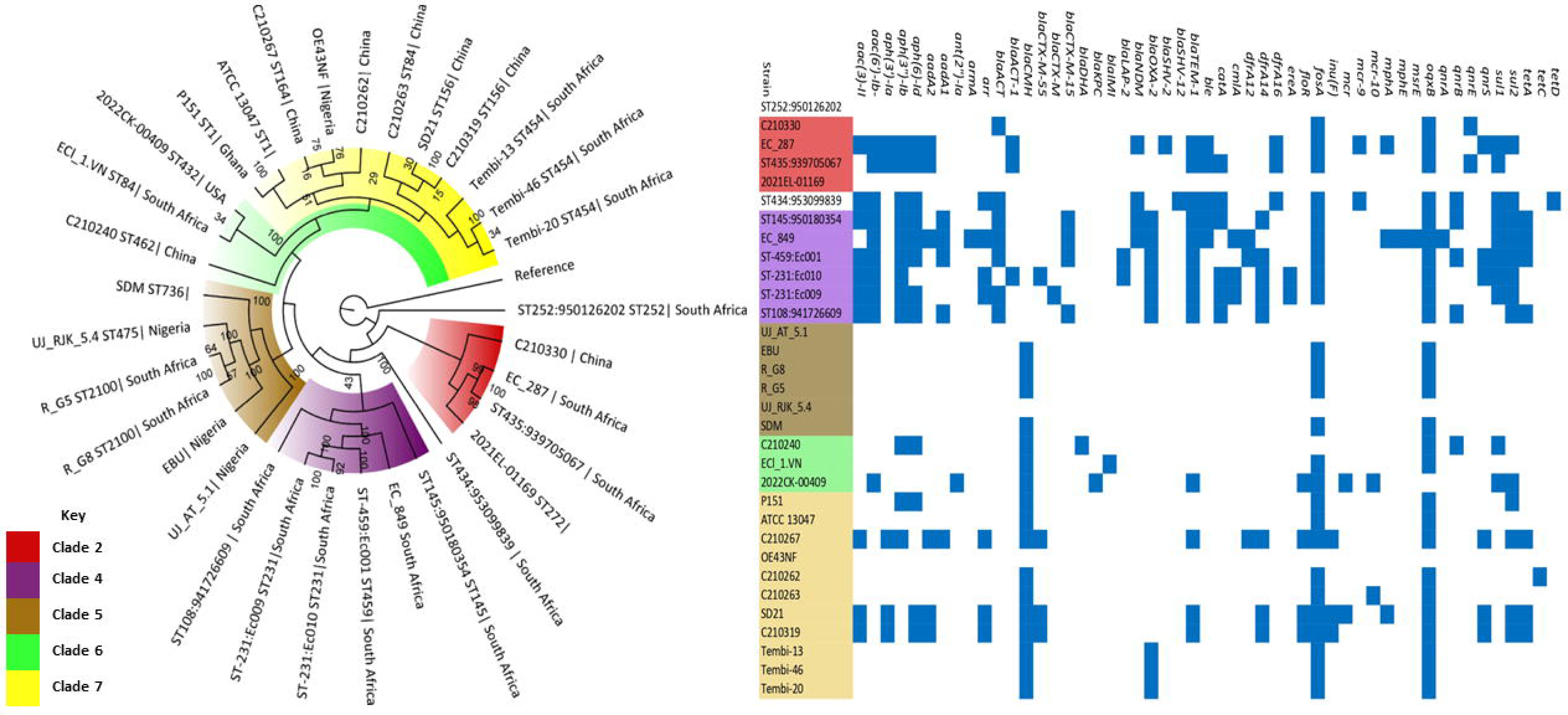
Global phylogenetic and resistome dynamics of *E. cloacae* complex isolates, collected from human samples. Each strain is represented by its strain identifier, MLST designation, and country of origin. Strains belonging to the same clade are highlighted with the same color on the branches. The resistome is depicted through blue and white blocks, representing the presence and absence of antibiotic resistance genes, respectively.

### 3.7 Epigenomics

Types I, II, and III Methyltransferases (Mtases) were detected in the sequenced isolates (n = 10). Among these, Type II Mtases were the most predominant, followed by type I Mtases. Conversely, type III Mtases were the least common, and type IV Mtases were not identified in any of the isolates (Figure 7).

**Figure 7.**
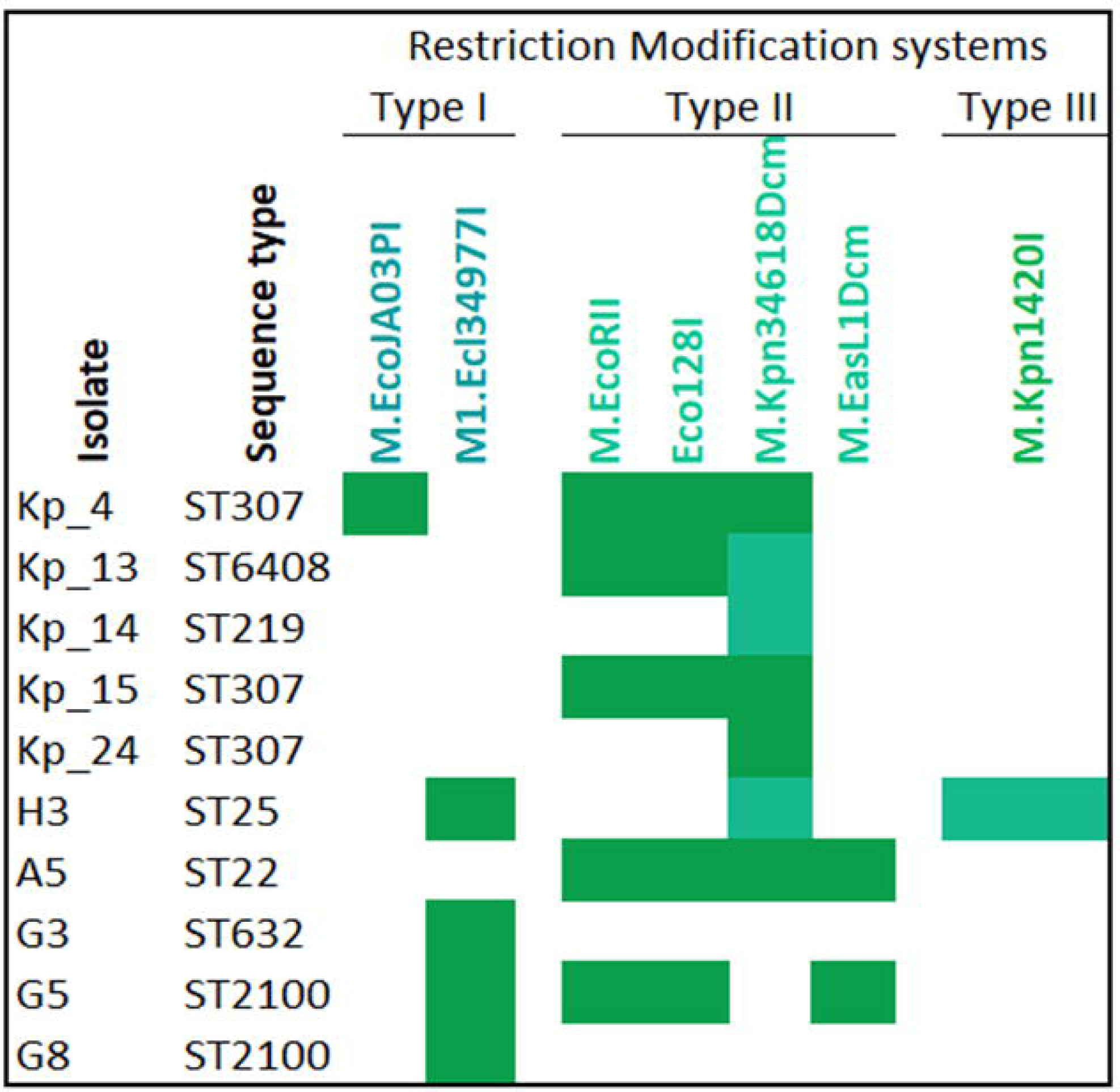
Distribution of Restriction-Modification (R-M) sites within the *Klebsiella pneumoniae* and *Enterobacter* species.

A single type III Mtase, *M.kpn1420I*, was located chromosomally within isolate H3, alongside a single Type I and II Mtase: *M1.Ec13497I* and *M.Kpn34618Dcm*, respectively. Each Mtase harboured by isolate H3 had its own unique recognition sequence. Lastly, isolate H3 was the only isolate that harboured three types of Mtase (Table S5).

A Type II restriction endonuclease (RE), *Eco128I*, was identified in five isolates: Kp_4, Kp_13, Kp_15, A5, and G5. Significantly, in each of these isolates, *Eco128I* was encoded by a plasmid. Interestingly, all four Type II Restriction-Modification Systems (RMS) identified in the isolates, including the RE, shared the same recognition sequence, CCWGG. The most common of these was M.*Kpn34618Dcm*, which was present in eight of the ten isolates. Notably, it was located chromosomally in the *K. pneumoniae* isolates Kp_4, Kp_13, Kp_14, Kp_24, and H3 while in isolates Kp_15, A5, and G5, it was plasmid encoded. This means that in isolate Kp_13, both a Type II RE and Mtase (M*.EcoRII* and *Eco128I*) were identified on a plasmid, alongside a type II Mtase (M*.Kpn34618Dcm*) within the chromosome. Notably, Type II Mtases were not identified in isolates G3 and G8.

The type II RMS adenine (Dam) Mtase was not found in any of the study isolates; only the type II RMS cytosine (Dcm) Mtase was present. Two Dcm Mtases were identified: M.*Kpn34618Dcm* and M.*EasL1Dcm*, with the latter only identified in isolate A5. A complete RMS consisting of REs, Mtases, and a specificity subunit was not found in any of the isolates, as no specificity subunits were identified during the analysis. Both an RE and Mtase were found in the five isolates encoding the Type II RE. These isolates, Kp_4, Kp_13, Kp_15, A5, and G5, further harboured the same type II Mtases, *EcoRII* and M*.Kpn34618Dcm*, with isolate A5 also harbouring an additional type II M*.EasL1Dcm*. The remaining five isolates only harboured MTases.

Two type I Mtases were detected: *M.EcoJA03PI* and *M1*.*Ec134977I*. They had distinct recognition sequences, GATGNNNNNCTG and GCCNNNNNGTT, respectively, and were both located chromosomally. *M1.Ec134997I* was present in four isolates: H3, G5, G3, and G8, while *M.EcoJA03PI* was only identified in isolate Kp_4.

As described in the methods, PacBio SMRT sequencing was only performed on five isolates: Kp_14, Kp_25, H3, A5, and G5. All isolates had m6A modifications that result in N6-methyladenine (6mA) modifications, with the GATC motif being identified in all isolates (Table S5). Moreover, the m4C modification, resulting in N4-methylcytosine (4mC), was also present in all isolates, with the VVNCYGVNYR motif identified in all cases.

### 3.8 Differential gene expression analysis

The analysis of differentially expressed genes (DEGs) was performed using HTSeq-DeSeq2 tool, and the data was visualized using SRPlot (seen in Figures 8-10). The DEGs’ data was further analysed on an Excel spreadsheet, wherein non-significant genes were filtered out. In the case of Kp_4, this filtering process reduced the number of DEGs from 4493 to 86, and this trend was observed across the remaining nine isolates.

**Figure 8.**
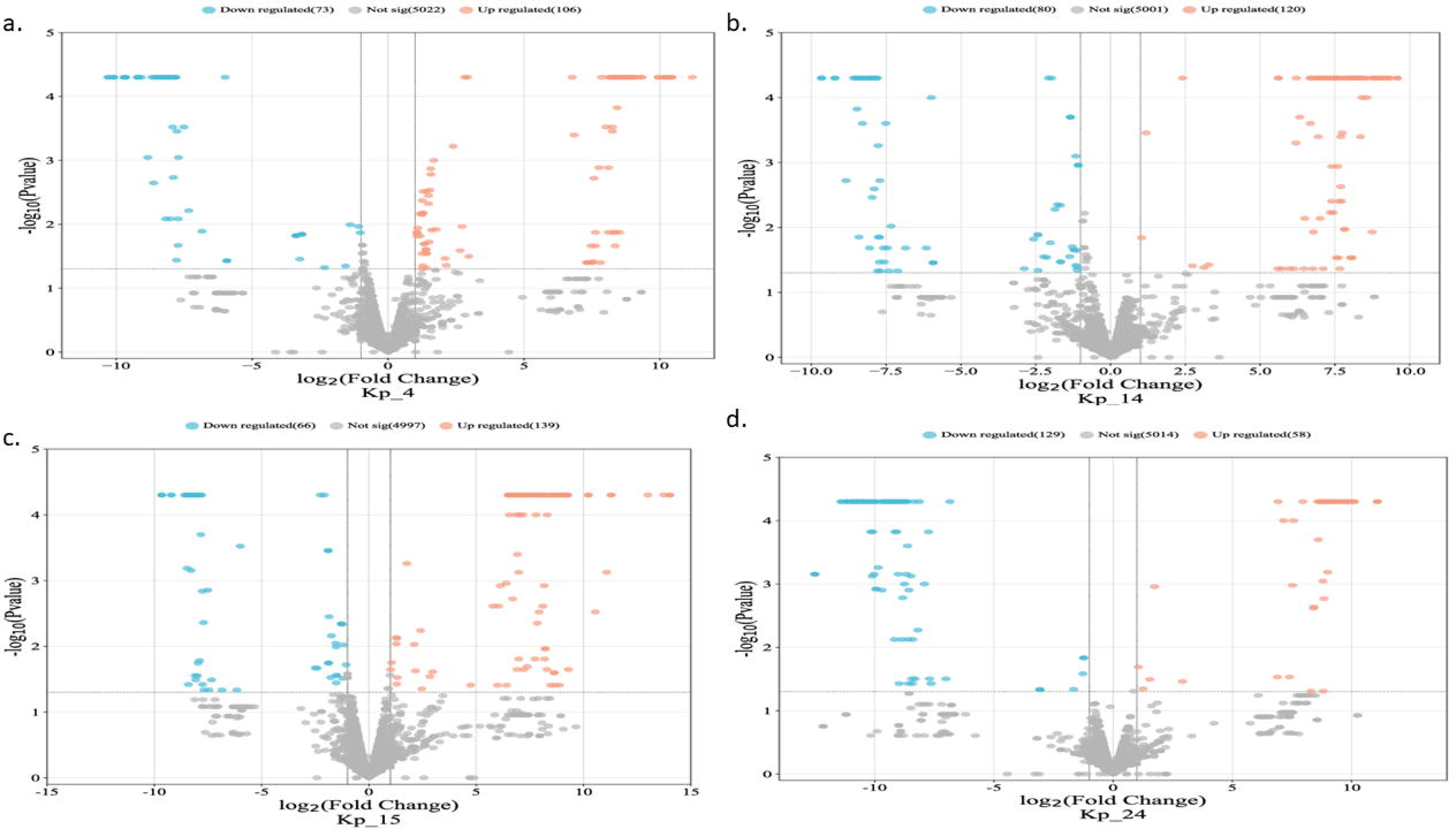
A volcano plot was used to compare the Differentially Expressed Genes (DEGs) between the carbapenem-resistant *K. pneumoniae* isolates and the susceptible Kp_13 isolate using *K. pneumoniae* as a reference genome. Each data point represents a gene, and its position was determined by the fold change (log2FC) and the statistical significance (log p-value). Orange dots represent upregulated genes, blue dots represent the downregulated genes, and grey dots represent the non-significant genes (P < 0.05, logFC > 0).

**Figure 9.**
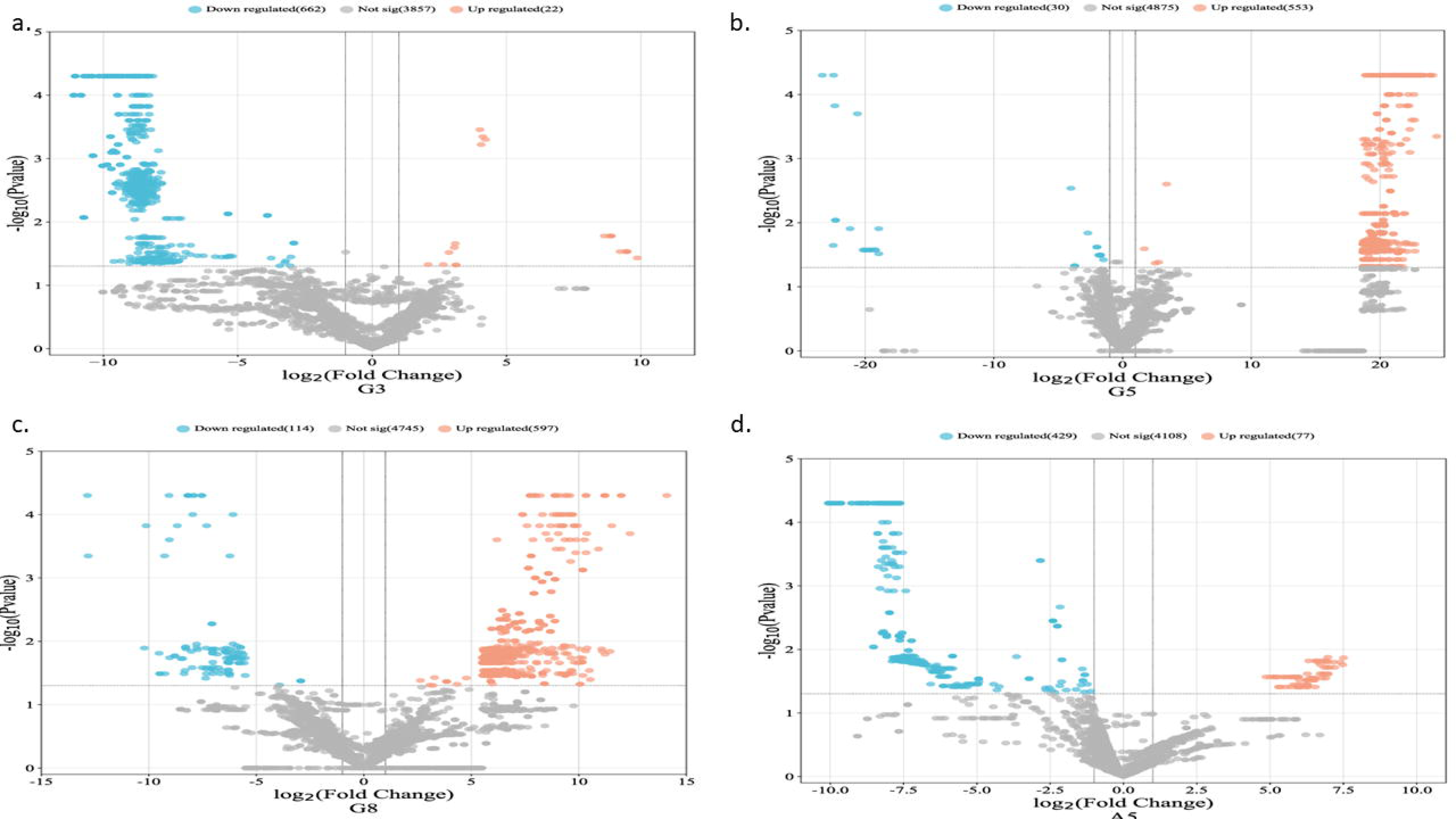
A volcano plot was used to compare the Differentially Expressed Genes (DEGs) between the colistin *Enterobacter sp.* isolates and the reference genome Each data point represents a gene, and its position was determined by the fold change (log2FC) and the statistical significance (log p-value). Orange dots represent upregulated genes, blue dots represent the downregulated genes, and grey dots represent the non-significant genes (P < 0.05, logFC > 0).

**Figure 10.**
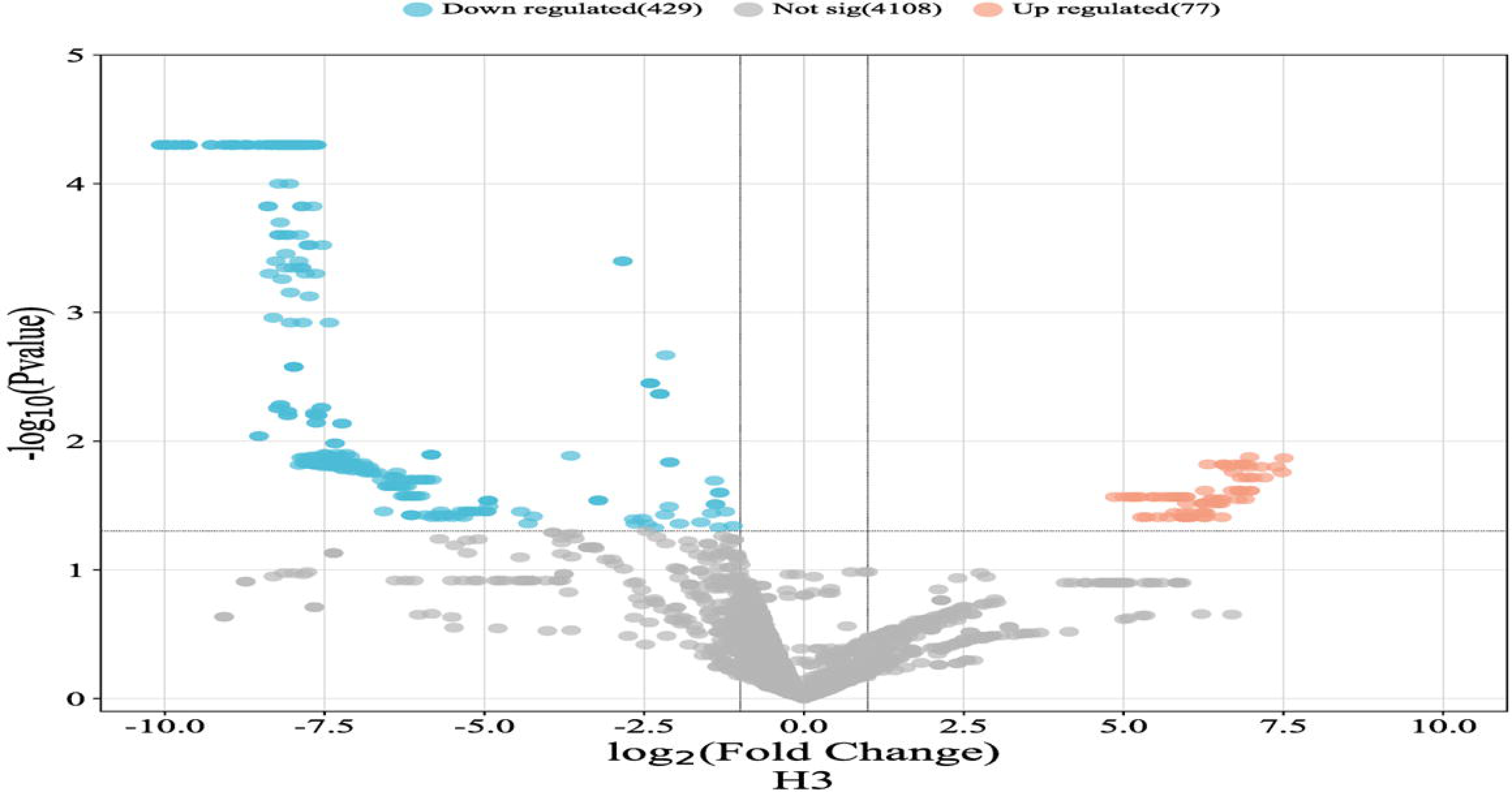
A volcano plot was used to compare the Differentially Expressed Genes (DEGs) between the colistin-resistant *K. pneumoniae* isolate, H3 and the susceptible Kp_13 isolate using *K. pneumoniae* as a reference genome. Each data point represents a gene, and its position was determined by the fold change (log2FC) and the statistical significance (log p-value). Orange dots represent upregulated genes, blue dots represent the downregulated genes, and grey dots represent the non-significant genes (P < 0.05, logFC > 0).

The patterns of DEGs were found to be similar in eight isolates (G5, G8, H3, Kp_4, Kp_14, Kp_15 and Kp_24), as seen in Table S6-S7, with capsular polysaccharide biosynthesis genes showing increased expression. This upregulation was seen in isolate Kp_14 and Kp_15. Moreover, changes were observed in the membrane area of the clinical isolates, including the downregulation of ion ABC-transporters in all *K. pneumoniae* isolates (Kp_4, Kp_14, Kp_15 and Kp_24).

Isolate G5, G8, Kp_14, and Kp_15 displayed increased expression of three ion-ABC transporters: an ATP-binding protein, permease protein, and a substrate-binding protein. Additionally, the ferric-ion transporter was upregulated in Kp_14 and Kp_15 isolates, while there was a downregulation of Iron (III) dicitrate transporter in Kp_14 and Kp_24 (Table S6). Isolate G5 had an upregulation of the ferric hydroxamate outer membrane receptor, FhuA.

The core metabolic functions also had differential expression; *sufAB,* responsible for iron-sulfur metabolism, showed increased expression in all *K. pneumoniae* isolates. On the other hand, cobalt-precorrin methyltransferase was downregulated in Kp_4, Kp_14, Kp_15 and Kp_24. The putative glycotransferase, involved in the biogenesis of natural products, was upregulated in all *K. pneumoniae* isolates; and in isolate G5, this protein was additionally upregulated along with an LPS core biosynthesis glycotransferase and an LPS core heptosyltransferase. Additionally, D-3 phosphoglycerase dehydrogenase had upregulation in Kp_4, Kp_14, Kp_15, and Kp_24. Lastly, the cellulase synthase was upregulated in Kp_14 and Kp_15 isolates, while a 3-oxoacyl-[acyl carrier protein (ACP)] synthase was upregulated in isolates G5 and G8.

In the *K. pneumoniae* isolates, seven transcriptional regulators were upregulated (Tables S6 and S7). Among these were a probable transcriptional regulator of MDR efflux pumps and a transcriptional regulator associated with rhamnose utilization, part of the AraC family, were upregulated in all *K. pneumoniae* isolates (Table S6). In isolate G5, four transcriptional regulators were upregulated (Table S7). One of these regulators belongs to the AcrR family, responsible for regulating the AcrAB-TolC MDR efflux system, was upregulated alongside H3. Additionally, the RND efflux pump regulator was also upregulated in isolate G5 along with isolate G8.

Components of the type 1 fimbriae were found to be upregulated in all *K. pneumoniae* isolates and in isolate G5. These components include the outer membrane usher protein, fimbrial protein *staA*, the fimbrial protein subunit precursor and the fimbrial chaperone.

## Discussion

The emergence of colistin- and carbapenem-resistant *K. pneumoniae* is a major concern owing to limited treatment options. Epidemiological data in South Africa shows an increased prevalence of carbapenemase-positive Gram-negative bacteria and a low prevalence of *mcr* genes within the public health sector^4,14,25,26^. However, there are carbapenem- and colistin-resistant isolates without any known resistance mechanism. This study, therefore, aimed to characterize novel colistin and carbapenem resistance mechanisms in clinical *K. pneumoniae* isolates from South Africa.

Four non-carbapenemase producing carbapenem-resistant *K. pneumoniae* and five non*-mcr* producing colistin-resistant Enterobacteriacae species were examined. Although the colistin-resistant isolates were identified by Microscan as *K. pnuemoniae*, only isolate H3 was confirmed to be *K. pneumoniae.* The remaining isolates were identified as *Enterobacter* species.

The Microscan analysis showed that the *Enterobacter* species had reduced susceptibility to β-lactams, β-lactams/β-lactamase inhibitors, as well as the first- and second-generation cephalosporins. The resistance mechanisms associated with these antibiotics involve β-lactamase activity and loss of porin activity ^27-29^. The *Enterobacter* species, G5, and G8 harboured *bla*_CMH_, which is the most common β-lactamase gene within the *Enterobacter* genus. Additionally, *bla*_ACT_ which is also commonly found in this genus ^30^, was present within A5 and G3. The *Enterobacter* species also harboured three other resistance genes: *fosA*, conferring resistance to Fosfomycin ^31^, *oqxAB*, conferring resistance to quinolones, tigecycline, nitrofurantoin, several detergents, and disinfects ^32^. No other resistance genes were identified. However, the phenotypic characterization of isolates revealed reduced susceptibility to ertapenem, meropenem, colistin and tobramycin. Resistance to these antibiotics can be mediated through changes in the outer membrane permeability, alteration of the lipopolysaccharide reducing porin activity and increased activity of efflux pumps ^11,33^.

The efflux pump inhibition assay showed that isolates G5, G8 and H3 had increased susceptibility to colistin in the presence of CCCP efflux pump inhibitor (EPI). The colistin BMD MIC value reduced 1-fold from 128 µg/mL to 64 µg/mL. This EPI has been shown to restore colistin susceptibility in intrinsic colistin resistant bacteria in some Enterobacteriacae isolates ^34,35^. Although colistin susceptibility was not fully restored in these isolates, the inhibition of efflux pump activity highlights their role in colistin resistance.

Colistin resistance has been previously linked to *mcr* activity ^36^, modification of the lipopolysaccharide (LPS) ^37^, overexpression of efflux pumps ^38^, and overproduction of capsular polysaccharide ^39-41^. Genomic analysis reveals that the colistin-resistant isolates (A5, G3, G5, G8 and H3) were sorely negative for *mcr* gene. However, H3 harboured a *mgrB* mutation that has been demonstrated to confer colistin resistance by regulating the LPS modification system^17^. The colistin resistant isolates, as revealed by RNA-seq analysis, showed a range of potential mechanisms for mediating resistance, including upregulation of efflux pumps, capsular polysaccharide biosynthesis, and putative glycosyltransferases. Common membrane alterations in colistin-resistant strains encompass the upregulation of MDR efflux pumps and capsular polysaccharide biosynthesis, which could potentially mediate colistin resistance. Moreover, the upregulation of the *fimH* and capsule genes, coupled with the presence of the *mrkA* virulence factor, might facilitate biofilm formation, thereby promoting antibiotic resistance ^42^.

The production of capsular polysaccharide was observed in isolates G5, G8 and H3. Previous studies have indicated that this activity acts as a protective barrier against cationic antimicrobial peptides like colistin ^39^. As a result, this reduces the interactions between colistin and the LPS, thereby mediating resistance. Putative glycosyltransferase, notably those encoded by *crrB* gene, has been shown to mediate the LPS outer membrane modification ^43^. The observed upregulation of putative glycosyltransferase in isolates G5 and H3 suggests a potential role in mediating LPS modifications. Telke *et al* (2019) previously reported that the overexpression of the acrAB-tolC efflux pump, regulated by *soxRS* in *E. cloacae* and *E. asburiae* isolates, resulted in colistin hetero-resistance ^44^. In our study, the efflux pump activity observed in G5, G8, and H3 was regulated by the *acrR*, as seen in Table S7. However, the remaining isolates, G3 and A5 did not display significant DEGs that could confer colistin resistance. The detailed DEG Tables for each isolate can be found in Table S8-16.

The *K. pneumoniae* isolates harboured a wide range of ARGs that confer resistance to various classes of antibiotics. These include aminoglyocsides (*acc(3)-IId, aac(6’)-Ib-cr, aadA2, aadA16, aph(3’)-Ia, aph(3”)-Ib, aph(6)-Id, armA, strAB),* cephalosporins *(bla*_CTX-M_), quinolones (*oqxA, oqxB*), fosfomycin (*fosA*), pencillins (*bla*_TEM_, *bla*_DHA_, *bla*_OXA_, *bla*_CMH_, *bla*_SHV_), sulfonamides (*sul1, sul2*), tetracyclines (*tetA*), and trimethoprim (*dfrA*). The genes contributed to the observed phenotypic resistance. Isolates Kp_4, Kp_15 and Kp_24 harboured mutations in *ompK35,* which confer resistance to carbapenems^11,45^.

DNA analysis revealed that all the carbapenem-resistant *K. pneumoniae* (Kp_4, Kp_14, Kp_15, and Kp_24) isolates harboured multiple β-lactamases and mutations within *ompK36* and *ompK37,* except for isolate Kp_14. The combination of porin mutations in *ompK* and β-lactamase activity contributes to carbapenem resistance. Additionally, these isolates exhibited upregulation of MDR efflux pumps ^46,47^.

RNA-seq analysis revealed that the carbapenem-resistant *K. pneumoniae* isolates had multiple mechanisms to confer resistance to carbapenems. These resistance mechanisms included the production of capsules, biofilm formation, and increased efflux activity. Interestingly, in isolates Kp_14 and Kp_15, capsule polysaccharide biosynthesis was coupled with the upregulation of cellulose synthase. This coupled upregulation has previously been demonstrated to facilitate biofilm formation ^46^. Furthermore, all the carbapenem-resistant isolates exhibited a wide variety of upregulated fimbriae products (Table S6). Additionally, the analysis revealed the upregulation of an *acrR* transcriptional regulator and a probable MDR transcriptional regulator protein, further highlighting the complex mechanisms at play in conferring carbapenem resistance.

In contrast, the EDTA and efflux pump inhibition analysis (Table 2) demonstrated that β-lactamase activity and efflux pumps played a role in carbapenem resistance in all isolates except Kp_24. Therefore, in this group, three distinct resistance mechanisms were observed, excluding Kp_24, where efflux pump activity did not contribute to carbapenem resistance.

Table 3 shows that majority of ARGs identified within the *K. pneumoniae* isolates were harboured on plasmids. IncFIB(K), IncFIB(K)/IncFII(K), and IncHIB harboured three or more resistance genes with the IncFIB(K)/IncFII(K) harbouring a remarkable nineteen resistance genes (Table S2). These plasmid replicons, IncF and IncH, are among the most observed types of replicons in Enterobacteriaceae, and they play a significant role in facilitating transmission of ARGs ^48-50^. Studies have shown that IncFIB and IncFII replicons are capable of accommodating and stably carrying a wide variety of ARGs ^51-53^. These accounts for the large number of resistance genes seen in the IncFIB(K)/IncFII(K) plasmid from this study. Furthermore, it was observed that the ARGs within these plasmids were often flanked by IS elements, particularly IS26, which is widely known to be associated with ARGs ^54,55^. This underscores the potential role of IS elements in these isolates in the dissemination of these MDR ARGs, thus facilitating the widespread of ARGs in South Africa. This may occur, through the transfer of ARGs between animal-derived and human derived pathogens.

Genomic analysis of the six *K. pneumoniae* isolates revealed that the isolates belonged to four sequence types. ST307 clone comprised three isolates, Kp4, Kp15 and Kp24, which had the same K- and O-serotypes, KL102 and O1/O2vO2. The K-antigen describes the type of capsular polysaccharide harboured by the *K. pneumoniae* isolates, and the O-antigen describes the lipopolysaccharide antigens ^56^. The KL102, previously known as KN2, has been widely identified in carbapenemase-producing *K. pneumoniae* isolates in Nigeria ^13^, USA ^57^ and Switzerland ^58^. These isolates were further shown to also harbour the O1/O2v2 serotype. However, in this study, despite the isolates harbouring the same sequence type and serotypes, the phylogenetic analysis of these isolates revealed an interesting pattern in their distribution and resistance profiles. The analysis included 81 *K. pneumoniae* isolates from five continents, with sequence types ST307, ST25, and ST219 being the most common. Within South Africa, the majority of *K. pneumoniae* isolates belonged to ST307, and five of the eight Clades were comprised of this sequence type. These five Clades had similar resistomes, thus highlighting the vertical and horizontal spread of this MDR clone and ARGs within South Africa. Furthermore, in Figure 3, the study isolates (Kp_4, Kp_13, Kp_14, Kp_15, and Kp_24) clustered alongside international *K. pneumoniae* isolates, underscoring their easy transmissibility and wide distribution.

The *Enterobacter* species included isolates A5, G3, G5, and G8, which were identified as *Enterobacter asburiae*, *Enterobacter bugandensis,* and two *Enterobacter cloacae* species, respectively. The sequence types of these isolates included ST22 (A5), ST632 (G3) and a novel ST2100 for both *E. cloacae* species (G5 and G8). Fortunately, these isolates carried only a limited number of resistance genes and lacked plasmids. Moreover, they clustered with other isolates that had similar resistance patterns. Specifically, *E. asburiae* A5 clustered within clade 4 (Figure 4) and clustered with a South African strain (E124_11) and a Chinese strain (C210176). Notably, this clade displayed a distinct resistome pattern compared to the other clades; a pattern consistent with the other *Enterobacter* species analysed in this study (Figures 5 and 6). This distinction might be attributed to the presence of different plasmids that potentially encode these ARGs. A more comprehensive phylogenetic analysis, which incorporates plasmid analysis of the included isolates, could shed light on the reasons behind this clustering pattern.

The study’s isolates were found to harbour a diverse array of restriction modification systems (RMS), including both restriction enzymes and methyltransferases. These RMS included Types I, II, and III RMS. Among these, the Type II *M*.*Kpn34618Dcm* was the most predominant and was identified in all *K. pneumoniae* isolates. Previous research, as reported by Chuckamnerd *et al.* 2022 and Ramaloko and Osei Sekyere (2022), has shown the common occurrence of this Mtase in *K. pneumoniae ^59,60^*. Moreover, it is typically found alongside *M.EcoRII*, a pattern noted by Ramaloko and Osei Sekyere (2022). In this study, it was observed that four of the seven isolates harbouring the Dcm Mtase also carried a plasmid-encoded *M.EcoRII*. Interestingly, the *E. asburiae* A5 isolate displayed a similar combination of these Mtases.

In contrast, among the ST307 *K. pneumoniae* isolates (n = 3), *M.Kpn34618Dcm* was the sole common Mtase. Contrary to Chuckamnerd *et al.* (2022) findings, there was no consistent pattern observed within the RMS in this study’s isolates ^59^. However, it’s noteworthy that all Type II Mtases, including the type II restriction endonuclease (RE), shared the same recognition sequence. This commonality facilitates the integration of plasmids encoding these Type II RMS into host bacteria, thereby enhancing the dissemination of virulence and resistance genes ^6^.

Only two of the three types of methylation i.e., N6-methyladenine (m6A) and N4-methylcytosine (m4C), were identified in the isolates that underwent PacBio SMRT sequencing (Kp_4, Kp_24, A5, G5, and H3). According to Militello *et al.* (2012), the methylation type N5-methylcytosine (5mC) DNA modification, is not commonly found ^61^. In this study, neither *K. pneumoniae* nor the *Enterobacter* species isolates encoded this type of methylation. However, m6A and m4C, representing an alternative form of cytosine methylation, were detected. It is noteworthy that only a small fraction of motif sites in the isolates remained non-methylated, as depicted in Table S5.

In addition to resistance mechanisms, the *K. pneumoniae* isolates also carried various virulence genes, making them highly equipped for pathogenesis. The isolates harboured nine types of virulence genes, including adhesion, biofilm formation, efflux pumps, immune evasion, iron uptake, regulation of capsule synthesis, and secretion systems. The presence of these virulence genes further underscores the necessity for effective infection control measures to prevent the spread of these highly virulent and drug resistant strains.

Virulence genes play a pivotal role in the pathogenesis of a pathogen, facilitating both host infection and, in this case, resistance to antibiotics ^62^. The transcriptomic analysis revealed an increase in certain transporters, such as those for carbohydrates, cysteine, and ferric ions. Cain *et al* (2018) explained that signs of stress in *K. pneumoniae* include the accumulation of compounds like cellulase, carbohydrates and metal ions in granules at the end of active growth ^63^. Thus, the upregulation of ion ABC-transporters, phosphotransferase system components, and ferric-ion transporters may indicate stress induced by antibiotic exposure in these *K. pneumoniae* isolates. Ramos *et al.* (2016) further demonstrated that intracellular regulation of iron metabolism assists bacteria in managing oxidative stress ^64^. Another indicator of stress is the upregulation of fimbriae genes, as observed by Cain *et al.* (2018) ^63^. The transcriptomic data indicated upregulation of type 1 fimbriae genes, potentially mediated by *fimH* virulence gene ^65^.

Unfortunately, due to financial restrictions, the study was unable to employ the CRISPR-Cas system to investigate these putative resistance mechanisms in the clinical isolates. However, the combination of whole genome sequencing, epigenomics, and transcriptomics proved valuable in characterizing these resistance mechanisms.

Given the increasing prevalence of colistin and carbapenem-resistant *K. pneumoniae* in South Africa and globally, surveillance studies are essential to monitor the epidemiology and antibiotic susceptibility patterns of these MDR strains. This study contributes significantly to our understanding of the mechanisms behind antibiotic resistance and virulence in both *K. pneumoniae* and *Enterobacter* species. It offers valuable insights into the genomic, epigenomic, and transcriptomic characterization of colistin and carbapenem resistance mechanisms in clinical *K. pneumoniae* and *Enterobacter* species. The findings underscore the importance of continuous monitoring of the epidemiology and evolution of these pathogens.

Understanding the genetic basis of antibiotic resistance and virulence in *K. pneumoniae* is crucial for developing effective strategies to control and manage infections caused by these MDR bacteria.

## Supporting information

Table S1: Phylogenomic data of strains included in phylogenetic analysis.

Table S2: Mobile genetic elements data of isolates and their association with antibiotic resistance genes.

Table S3: Antibiotic resistance genes analysis of K. pneumoniae isolates.

Table S4: Virulome data of isolates and their association with mobile genetic elements.

Table S5: Restriction modification systems within isolates.

Table S6: Summarized differential gene expression data of carbapenem resistant isolates.

Table S7: Summarized differential gene expression data of colistin resistant isolates.

Table S8: Kp_4 differential gene expression data

Table S9: Kp_14 differential gene expression data

Table S10: Kp_15 differential gene expression data

Table S11 Kp_24 differential gene expression data

Table S12: A5 differential gene expression data

Table S13: G3 differential gene expression data

Table S14: G5 differential gene expression data

Table S15: G8 differential gene expression data

Table S16: H3 differential gene expression data

## Acknowledgements

This work is based on the research supported wholly/in part by the National Research Foundation of South Africa under grant number: 131013.

## Transparency declaration

none

## Author contributions

MM undertook laboratory work and manuscript drafting; NMM was a co-supervisor to the study and assisted with funding; BF was a co-supervisor to the study and assisted in reviewing of the manuscript; JOS designed and supervised the study and reviewed the manuscript, as well as assisted with analysis of the data.

## Supplemental files

**Table S1:** Phylogenomic data of strains included in phylogenetic analysis.

**Table S2:** Mobile genetic elements data of isolates and their association with antibiotic resistance genes.

**Table S3:** Antibiotic resistance genes analysis of K. pneumoniae isolates.

**Table S4:** Virulome data of isolates and their association with mobile genetic elements.

**Table S5:** Restriction modification systems within isolates.

**Table S6:** Summarized differential gene expression data of carbapenem resistant isolates.

**Table S7:** Summarized differential gene expression data of colistin resistant isolates.

**Table S8:** Kp_4 differential gene expression data

**Table S9:** Kp_14 differential gene expression data

**Table S10:** Kp_15 differential gene expression data

**Table S11** Kp_24 differential gene expression data

**Table S12:** A5 differential gene expression data

**Table S13:** G3 differential gene expression data

**Table S14:** G5 differential gene expression data

**Table S15:** G8 differential gene expression data

**Table S16:** H3 differential gene expression data

